# Metabolomic and Transcriptomic rewiring between virus resistance and susceptibility in *Ostreococcus mediterraneus*

**DOI:** 10.64898/2026.02.10.705097

**Authors:** Amandin James, Rémy Marcellin-Gros, Sheree Yau, Didier Stien, Gwenael Piganeau

## Abstract

The genus *Ostreococcus*, comprising picoeukaryotic unicellular algae, plays a key role in many coastal marine ecosystems. These phytoplankton are infected by lytic double-stranded DNA prasinoviruses against whom they have the capacity to reliably evolve stable antiviral resistance. Regulation of virus resistance and susceptibility is hypothesized to arise through a phenotypic switch, as resistant cell lines can be isolated from susceptible lines after viral exposure and *vice versa*. To elucidate the molecular mechanisms underlying this resistance, we integrated untargeted metabolomic analyses and transcriptomics in virus-resistant and virus-susceptible lines of *O. mediterraneus*. Transcriptomic analyses corroborated previous findings in *O. tauri*, revealing that most gene expression changes were concentrated on a single chromosome. Specifically, a ∼530 kb region was over-transcribed in virus-resistant lines, while a distinct ∼110 kb region was over-expressed in virus-susceptible lines. Comparative metabolomics identified several oxidized galactolipids and oxidized sterols as biomarkers of the susceptible phenotype, while fewer biomarkers were identified for the resistant phenotype. By integrating transcriptomic and metabolomic signatures—focusing on the expression of genes within biosynthetic pathways linked to these metabolite biomarkers—we uncovered candidate molecular mechanisms underlying the cellular physiology of susceptible *versus* resistant phenotypes.

**Authors’ Summary:** Viruses are the most abundant biological entities in the ocean, shaping the dynamics of phytoplankton communities that underpin marine food webs. *Ostreococcus*, one of the smallest photosynthetic eukaryotes, is frequently infected by highly abundant prasinoviruses in its natural environment. In this study, we aimed to understand the physiological bases of virus-resistant and susceptible *Ostreococcus mediterraneus* lines. We compared gene expression and untargeted metabolite profiles between a resistant line and a susceptible line derived from it. The metabolic profiles were much more variable between replicates than the transcriptomic profiles and the most differentially expressed genes did not include those involved in the biosynthetic pathways of the metabolite biomarkers identified. We estimated the congruence between the metabolomes and transcriptomes as the percent of relative expression changes fitting to the relative change in the metabolite. This study provides evidence of the subtle links between gene expression and metabolomic signatures and the importance of integrating multiple levels of cellular processes.

## Introduction

Phytoplankton play a key role in marine ecosystems by sustaining primary production and contributing significantly to biogeochemical cycles through carbon fixation and oxygen production [1]. Among them, picophytoplankton of the genus *Ostreococcus* (Mamiellophyceae), which was first isolated from the Thau Lagoon in the Mediterranean Sea, France [2], represents one of the smallest known free-living eukaryotes, both in terms of cell (< 1 µm) and genome size [3]. Whole genome sequencing approaches have revealed that species within the *Ostreococcus* genus are more divergent than within the *Saccharomyces* genus [4], while metabarcoding surveys confirmed their worldwide distribution in coastal marine environments [5]. Concurrently, environmental surveys have demonstrated the ubiquity and high abundance of nucleocytoplasmic large DNA viruses (NCLDVs, *Nucleocytoviricota* phylum) in marine environments [6]. This includes the genus *Prasinovirus*, which specifically infect members of the Mamiellophyceae lineage such as *Ostreococcus* [7], *Bathycoccus* [8] or *Micromonas* [9]. Phytoplankton infecting viruses act as major ecological players in marine ecosystems, notably through the “viral shunt”, which redirects organic carbon and nutrients away from heterotrophic grazers into dissolved organic carbon [10].

Previous studies in *Ostreococcus* have emphasised the involvement of specific genomic regions, notably the Small Outlier Chromosome (SOC) in the evolution of resistance to prasinoviruses. This SOC is characterised by a low GC content, a paucity of orthologous genes, a higher density of transposable elements [3,11] and marked size variation among natural populations [12,13]. In *O. tauri*, differential gene expression analysis comparing resistant and susceptible strains revealed a significant overrepresentation of genes encoded on the SOC, supporting the hypothesis that this chromosome may play a central role in antiviral resistance [14].

A switch between resistant and susceptible cells, akin to a bet-hedging strategy, has been suggested to explain the dynamics of *Ostreococcus mediterraneus* strain RCC2590 host cells and it prasinovirus, OmV2 [11]. In this model, a subpopulation of cells spontaneously transitions to a virus-resistant phenotype within a predominantly susceptible population. Following viral infection and lysis of susceptible cells, the resistant subpopulation survives, as it remains unaffected by subsequent viral exposure. Conversely, within a virus-resistant population, virus-susceptible cells were able to be isolated, suggesting that a subpopulation of cells reverted to a virus-susceptible state [11]. This model allows the coexistence of a predominantly resistant population with infectious viruses as the subpopulation of susceptible cells enable continued viral replication. However, the molecular mechanisms involved in the susceptible and resistant phenotypes remain to be elucidated.

To date, metabolomic profiles associated with virus resistance remain uncharacterized in the Mamiellophyceae. Here, we address this gap by investigating the transcriptomic and metabolomic signatures of resistant and susceptible lines of *Ostreococcus mediterraneus* RCC2590. Metabolomic approaches have previously revealed extensive virus-induced metabolic rewiring in the model haptophyte *Gephyrocapsa huxleyi* (formerly *Emiliania huxleyi*), infected by E. huxleyi virus (EhV) (species *Coccolithovirus huxleyi*), another virus of the *Nucleocytoviricota* phylum. These analyses revealed profound reprogramming of host metabolism to support viral replication [15], including the virus-induced remodelling of sphingolipid metabolism to produce virus-specific glycosphingolipids (vGSLs), which are essential for virion assembly and release [16–18]. This metabolic remodelling is accompanied by additional host responses, such as mitochondrial dysfunction and calcium influx [19]. Beyond infection markers, distinct classes of glycosphingolipids further discriminate *G. huxleyi* host phenotypes. Notably, resistance-specific GSLs (resGSLs), characterised by the long-chain base d19:4, are overrepresented in resistant strains [20]. These findings suggest a central role of glycosphingolipid diversity in the molecular arms race between *G. huxleyi* and its virus, and the power of metabolomics to better understand the molecular bases of phytoplankton-virus interactions.

In *O. mediterraneus*, we first analysed transcriptomes and metabolomes independently to identify genes and compounds associated with viral susceptibility and resistance. We then integrated these datasets using a pathway-centred approach, and focused on the biosynthetic routes underlying metabolomic biomarker production. This enabled us to uncover the transcriptional programs supporting metabolic reconfigurations in susceptible and resistant phenotypes. Integrating transcriptomics and metabolomics allowed us to improve our systems-level understanding of the metabolic and transcriptional reprogramming associated with viral susceptibility and resistance.

## Results

One resistant (hereafter R) and one susceptible (hereafter S) line of *O. mediterraneus* were sampled over 24-hour day-night cycle (Figure S1). The R line was derived from a single-cell subculture of *Ostreococcus mediterraneus* RCC2590, which was free of viruses. The S line was previously isolated through experimental evolution of the R line using a single-cell dilution method, resulting in a clonal line that had become susceptible to the virus OmV2 (corresponding to line S1 in [11]). Transcriptomes were sequenced in triplicate at four different time points corresponding to distinct phases of the cell cycle; *t_1_* corresponds to early growth phase and initiation of photosynthetic activity, *t_2_* and *t_3_* mark the beginning and the end of the cell division cycle, respectively, and *t_4_* corresponds to the end of the dark phase (Figure S1). Along these four time points, over 93% of the genes were transcribed (7641 of the 8171) to some extent (≥ 10 aligned reads in at least one library) in both phenotypes. This observation aligns with previous transcriptome datasets from other *O. mediterraneus* strains (RCC2572, RCC1621, RCC2596, RCC2573, RCC2593, RCC1108) generated by The Marine Microbial Eukaryotic Transcriptome Sequencing Project (MMETSP) initiative [21]. Additionally, transcriptome data from the same strain RCC2590 [11] reported ∼7,500 gene transcripts, supporting the conclusion that a very high proportion of genes are expressed in this species.

### Transcriptomes are mainly shaped by temporal dynamics

Principal component analysis (PCA) of the transcriptomic dataset revealed that the first two axes together accounted for around 70% of the transcript variation among the 12 resistant and 12 susceptible samples, primarily grouping them by sampling time (Figure 1A). The first principal component (PC1), which explained 42.2% of the variance, separated samples into two temporal clusters: *t_1_* and *t_4_* (early light and late dark phases, respectively, separated by 4 hours in the day-night cycle) were positioned on the right side of PC1, while *t_2_* and *t_3_* (late light and early dark phases, collected 4 hours apart) were on the left. Although each time point was transcriptionally distinct, this distribution indicated that transcriptomic profiles were more similar between sample points separated by 4 hours over the 24-hour cycle, rather than transcriptomes separated by 8 hours. The second principal component (PC2) explained 29.5% of the variance, separating samples according to the light-dark phases. This supports a strong diel imprint on global gene expression, consistent with the synchronisation of the *O. mediterraneus* cell cycle with the day-night cycle, mirroring patterns observed in *O. tauri* [22,23]. PC1 and PC2 organised the samples in a circular arrangement reflecting the temporal order of sampling across the 24-hour cycle, with each time point forming a distinct cluster. Phenotype-specific transcriptional differences were captured along the fourth principal component (PC4), which accounted for 8.2% of the variance (Figure 1B). This separation indicates a stable, phenotype-specific transcriptional signature across all time points. Overall, this PCA highlights the interplay between temporal regulation, diel cycling, and phenotype in shaping the global transcriptome, thereby justifying the subsequent timepoint-by-timepoint differential transcriptome analyses.

**Figure 1.**
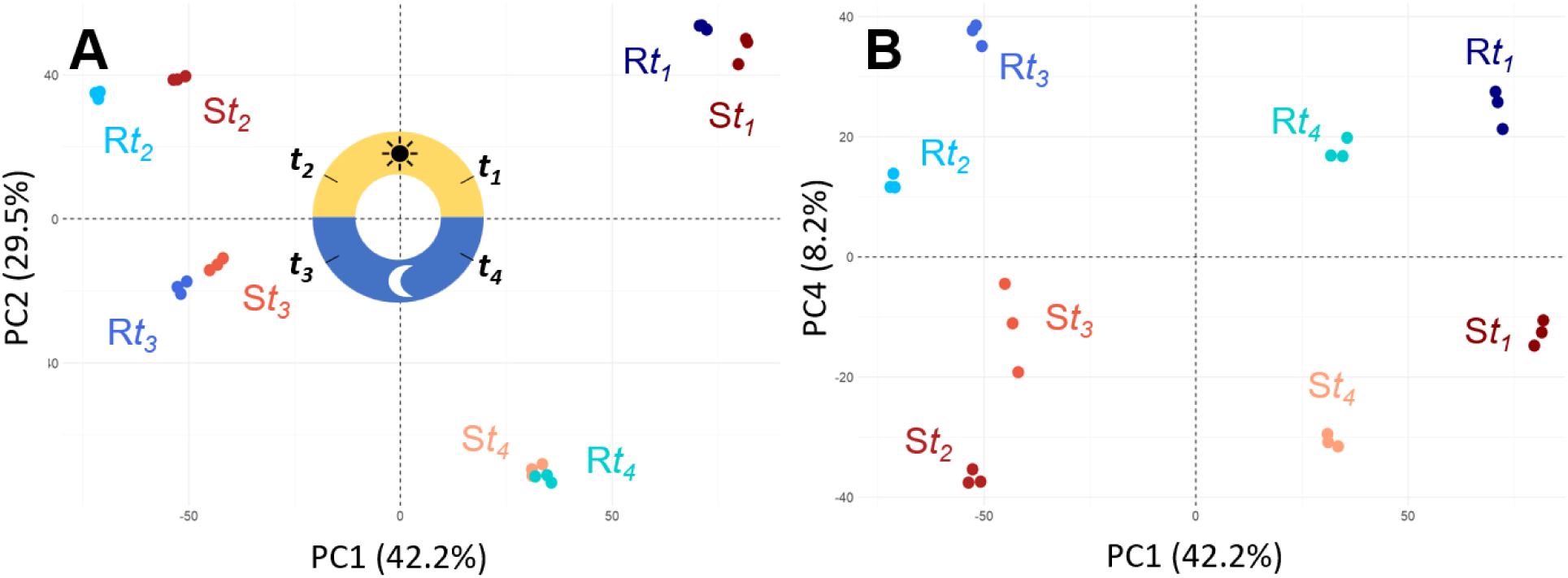
Principal component analysis of the normalized transcriptomes of susceptible (S) and resistant (R) *Ostreococcus mediterraneus* lines at sampling time points *t_1_*, *t_2_*, *t_3_*, and *t_4_*. Each point represents one sample. **A)** PC1 and PC2, **B)** PC1 and PC4.

### Most differentially transcribed genes are located on one chromosome with a bipartite structure

Differential gene transcription analysis comparing R and S lines at each corresponding sampling time point revealed 477 significantly differentially expressed genes (DEGs) with a fourfold difference in gene expression (|log_2_ fold| ≥ 2, *p*_adj_ < 0.01) at at least one time point. The highest variation in gene expression between the R and S lines occurred at the first time point, with 366 DEGs, compared to 248, 244, and 245 DEGs at *t_2_*, *t_3_*, and *t_4_*, respectively. Notably, only 184 genes (184/477=38%) remained consistently differentially expressed across all time points (Figure 2A).

**Figure 2.**
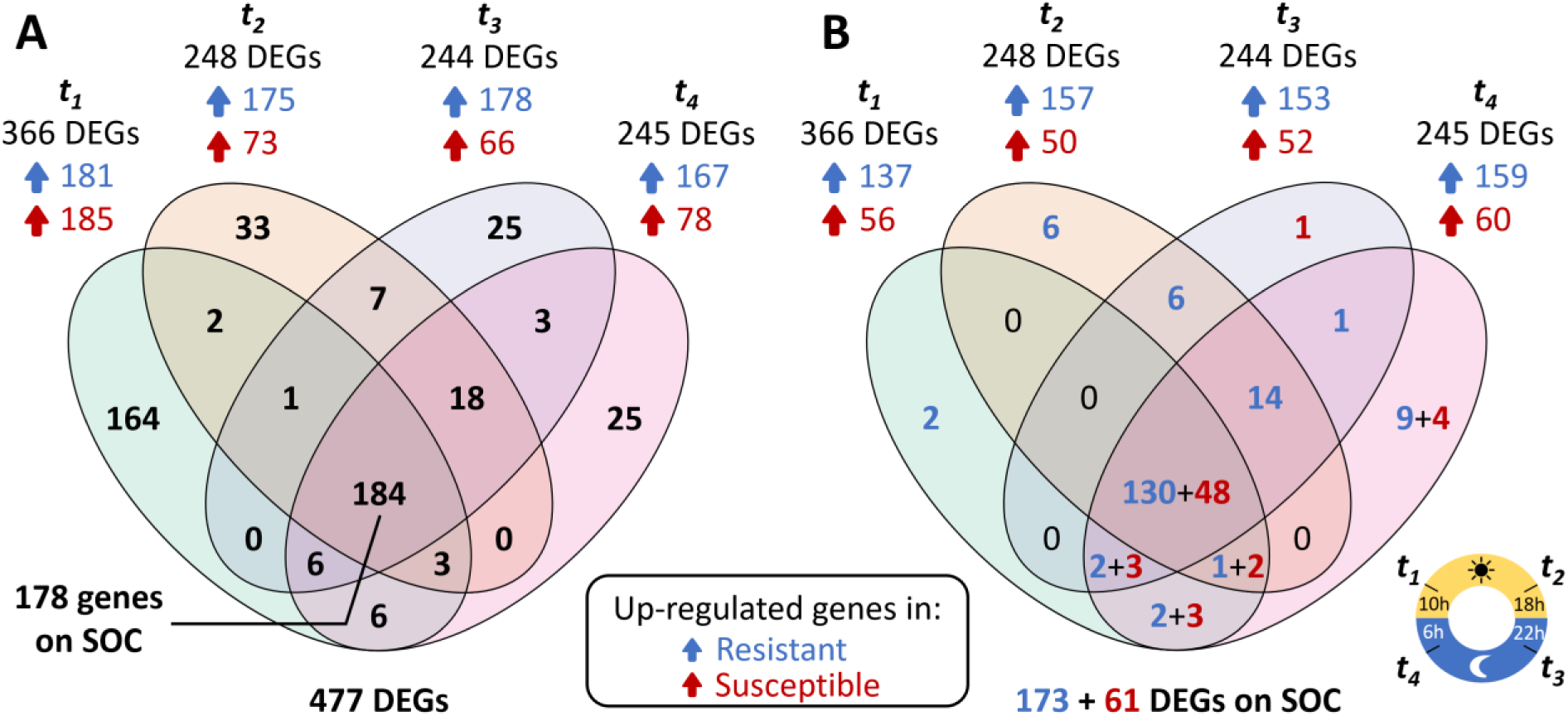
Number of Differentially Expressed Genes (DEGs) between resistant (R) and susceptible (S) lines in a Venn diagram. **A**) Total DEGs between the R (blue arrows) and S (red arrows) lines. **B**) DEGs on the small outlier chromosome (SOC). Each ellipse represents a time point of analysis, the values inside the ellipses and overlapping areas indicate the number of differentially expressed genes unique to or shared among the different time points.

The 477 DEGs showed a highly uneven distribution across the 20 genomic chromosomes of *Ostreococcus mediterraneus*. Strikingly, the small outlier chromosome (SOC) harboured the majority of DEGs, with 234 genes (Figure 2B). SOC-associated DEGs also displayed significantly higher log_2_ fold change (LFC) values compared to those on other chromosomes (Figure 3A, Figure S2), suggesting amplified transcriptional responses for genes on this chromosome. Of the 184 genes consistently differentially expressed across all four time points, 95% (178/184) were located on the SOC, representing 76% of all SOC DEGs (178/234, Figure S3). Among the 322 annotated genes on the SOC, 270 were transcribed (≥ 10 reads in at least one library), and 234 of these (87%) were differentially expressed at least once. This indicates that almost all transcribed SOC genes (87%) contribute to the observed transcriptional differences between resistant and susceptible lines.

**Figure 3.**
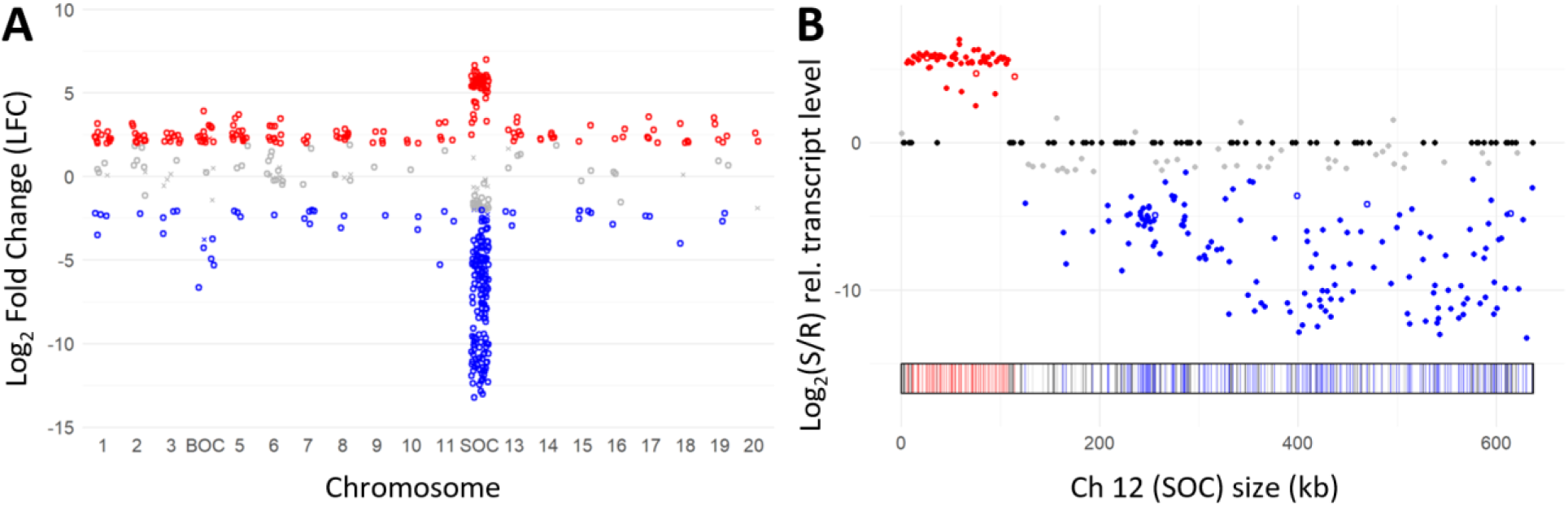
Differential gene expression between and within chromosomes. **A)** Distribution of log_2_ fold change (LFC) values across chromosomes for 477 differentially expressed genes (DEGs) at *t_1_*. Genes overexpressed (|LFC| ≥ 2) in the S line are shown in red, those overexpressed in R line in blue, and genes below this LFC are shown in grey. Significant values are shown as open circles and non-significant ones as crosses. **B)** Bipartite distribution of gene expression on the SOC at *t_1_*. Genes overexpressed in the S line are shown in red, those overexpressed in the R line in blue, not differentially expressed genes in grey, and non-expressed genes in black. Filled dots represent genes with “on/off” expression pattern, while empty dots indicate genes that expressed in S and R lines.

The chromosomal distribution of DEGs on the SOC revealed a bipartite organization. A ∼110 kbp region contained nearly all 61 genes overexpressed in the S line, whereas the adjacent ∼530 kbp region carried the 173 genes overexpressed in the R line (Figure 3B, Figure S2). Notably, this 530 kb region was transcriptionally silent in the S line, with gene expression either absent or minimal (< 10 reads). Based on previous work demonstrating an end deletion in the S line [11], we expected at least seven genes located at the right-hand end of the SOC (Figure 3B, Figure S2) to lack transcription, which was indeed the case. Among the 322 SOC genes, 62 showed no detectable transcription across all time points, suggesting potential silencing or deletion. Strikingly, over half of the SOC DEGs (122/234, *p*_adj_ < 0.01, |LFC| ≥ 2) exhibited an “on/off” transcription pattern, being fully repressed in one phenotype and actively expressed in the other. These findings indicate that the low transcription of most SOC genes in the S line results from transcriptional silencing, whereas their high transcription in the R line likely reflects active up-regulation.

### Functional analyses of DEGs

#### DEGs on standard chromosomes

Functional analysis of DEGs was performed using the Gene Ontology (GO) and InterPro domain (IPR) databases to characterize differences in metabolic and cellular processes between phenotypes (Table S1 to S4). Among the 243 DEGs not located on the SOC, of which 52% (126) had a GO annotation, and 70% of these DEGs were overexpressed in the S line.

When examining the susceptible line, several GO terms were significantly enriched (*p*_value_ < 0.05, Bonferroni correction) among overexpressed DEGs at *t_1_*, including functions related to ribosome assembly, transcription, translation and cellular metabolism. Specifically, these included the preribosome term (GO:0030684) and the nucleus term (GO:0005634). The nucleic acid binding term (GO:0003676) was also enriched. The organic cyclic compound binding (GO:0097159) and heterocyclic compound binding (GO:1901363) terms suggested a broader role in cellular metabolism and signalling. Several InterPro domains overrepresented in S lines were associated with RNA helicase functions, which are involved in RNA processing. These included the ATP-dependent RNA helicase DEAD-box (IPR000629); the RNA helicase, DEAD-box type, Q motif (IPR014014) domain; the DEAD/DEAH box helicase domain (IPR011545); the helicase, C-terminal (IPR001650); and the helicase superfamily 1/2, ATP-binding domain (IPR014001). No significant GO term enrichment was detected for overexpressed DEGs in S lines on standard chromosomes at timepoints *t_2_* and *t_4_*. At *t_3_*, an enrichment of geranylgeranyl reductase activity (GO:0045550) was identified (Table S1 to S2). This enzyme has a conserved role in the reduction of geranylgeranyl diphosphate, catalysing its stepwise conversion into a phytyl group, an essential precursor for chlorophylls, tocopherols, and phylloquinones.

In the resistant line, GO term enrichment among overexpressed genes was observed only at timepoints *t_3_* and *t_4_*. At *t_3_*, there was an overexpression of genes associated with nitrogen metabolism and processing. These included the nitrate metabolic process (GO:0042126) and nitrate assimilation (GO:0042128), nitrogen cycle metabolic process (GO:0071941), the reactive nitrogen species metabolic process (GO:2001057), and nitrogen compound transport (GO:0071705). Furthermore, the enrichment of molybdenum ion binding (GO:0030151) may also be linked to nitrogen metabolism, as molybdenum is a cofactor for enzymes like nitrate reductase involved in nitrate assimilation. At *t_4_*, there was an enrichment in the overexpressed genes associated with ion transport in the resistant phenotype, such as cellular potassium ion transport (GO:0071804) and potassium ion transmembrane transport (GO:0071805). There was also an enrichment of oxysterol-binding proteins (IPR018494 and IPR000648) that are key lipid transfer factors that regulate membrane lipid composition and function [24]. Finally, the enrichment in HTH CenpB-type DNA-binding domain (IPR006600) suggests a role in transcriptional regulation (Table S3 and S4).

In summary, DEGs between S and R phenotypes located on the standard chromosomes displayed marked temporal variation across the four time points. In susceptible cultures, overrepresented GO terms among overexpressed genes were predominantly associated with nuclear processes and RNA metabolism (*t_1_*), as well as a transient modulation of pathways related to isoprenoid metabolism or associated regulatory mechanisms (*t_3_*). Conversely, in resistant cultures, overrepresented GO terms among overexpressed genes were mainly nitrogen metabolism and processing (*t_3_*) and ion transport, particularly potassium ion homeostasis (*t_4_*).

#### DEGs on the SOC

Among the 234 DEGs on the SOC, only 17 had a GO annotation, 193, 207, 205, and 219 genes were identified as differentially expressed at *t_1_*, *t_2_*, *t_3_*, and *t_4_*, respectively. Of these, only 56, 50, 52, and 60 genes were specifically overexpressed in the S line at the corresponding time points consistent with little variation in gene expression over time (Figure 2B). In contrast, a substantially larger proportion (∼74%) of SOC DEGs were overexpressed in the R line. Importantly, all SOC DEGs were exclusively associated with either the S or the R phenotype, with no overlap between phenotypes.

The SOC genes from the S line exhibited enrichment for GO terms related to sulfotransferase activity (GO:0008146) and transferase activity, transferring sulphur-containing groups (GO:0016782), observed across all four time points, Additionally, the overexpression of genes with mannosyltransferase 1, CMT1 (IPR021047) and nucleotide-diphospho-sugar transferases (IPR029044) domains suggested their involvement in glycosylation-related processes. Transposon-associated domains, including the Harbinger transposase-derived nuclease domain (IPR027806) and PiggyBac transposable element-derived protein (IPR029526), were transcribed at *t_1_* and *t_4_* respectively.

The remaining 137, 157, 153, and 159 DEGs were specifically overexpressed in the R line at *t_1_*, *t_2_*, *t_3_*, and *t_4_*, respectively. In the R line, DEGs were consistently associated with GO terms related to DNA metabolism and dynamics were consistently identified across all time points, including DNA integration (GO:0015074) and DNA metabolic process (GO:0006259), which are involved in DNA recombination, repair, and regulation. The presence of glycosyltransferase groups (GO:0016757) at each time suggested enhanced glycosylation. At *t_1_* and *t_4_*, the enrichment of nucleic acid binding (GO:0003676) indicated interactions with nucleic acids. These results suggest elevated transposable element activity in the R line. The specific Interpro terms associated to these GO terms are detailed in Table S5 and S6. Additional domains, such as the Integrin alpha beta-propeller (IPR013519) and FG-GAP repeat (IPR013517) were also identified. These domains are typically associated with integrins, well-characterised cell adhesion molecules that mediate cell-extracellular matrix interactions and cell-cell communication (Table S5 and S6).

#### Genes systematically differentially expressed across time

Among the 477 DEGs identified across all time points, 184 were systematically differentially expressed at *t_1_*, *t_2_*, *t_3_* and *t_4_*. Strikingly, only 6 of these genes (∼3%) were located on the standard chromosomes, corresponding to merely ∼2% of the 243 DEGs mapped to these chromosomes. In contrast, 178 (∼97%) were located on the SOC, representing 76% of the SOC DEGs (178 out of 234). Of these 178 SOC systematically DEGs on the SOC, 48 genes were overexpressed in the S line, while 130 were overexpressed in the R line. GO terms associated with these DEGs were largely consistent with those identified for the full set of 234 DEGs (Table S7 and S8). However, notable differences are detailed below.

In the S line, the GO terms “integral component of membrane” and “intrinsic component of membrane”, were enriched among DEGs on the SOC overexpressed across all four time points. The broader term intrinsic component of membrane (GO:0031224) includes proteins anchored *via* transmembrane regions or covalent modifications, such as glycosylphosphatidylinositol anchors. Such proteins contribute to membrane integrity, protein-protein interactions, and cellular communication. The enrichment of these terms among systematically overexpressed genes on the S line suggests a potential role of membrane-associated processes in the regulation of cellular responses to viral infection or membrane competence during infection.

Conversely, in the R line overexpressed DEGs, the GO term nucleic acid binding (GO:0003676) observed at *t_1_* and *t_4_* in the broader set, was absent among the 178 SOC systematically DEGs across the four time points.

#### Biological functions of systematically DEGs on the SOC

To characterise the potential biological functions of the 178 SOC DEGs across all four time points, we manually assigned each gene to functional categories based on domain annotations (Table S9). Among the 48 DEGs overexpressed in the S line, the most represented categories included glycosyltransferases, signal peptides, methyltransferases, and endonucleases (9, 4, 3 and 4 genes respectively). In contrast, among the 130 genes overexpressed in the R line, the dominant functional categories were glycosyltransferases, signal peptides, reverse transcriptases and endonucleases (11, 13, 12 and 9 genes respectively). Additional functional groups specific to the resistant line included epimerases, hydrolases, peptidases, ribonucleases, and transposases (3, 4, 2, 2 and 2 respectively), whereas only one transposase and no hydrolase, epimerase, or peptidase were found in the DEG overexpressed in the S line. A substantial proportion of DEGs lacked functional annotations and were classified as unknown, representing 67 genes (52%) in the R line and 22 genes (46%) in the S line (Figure 4).

**Figure 4.**
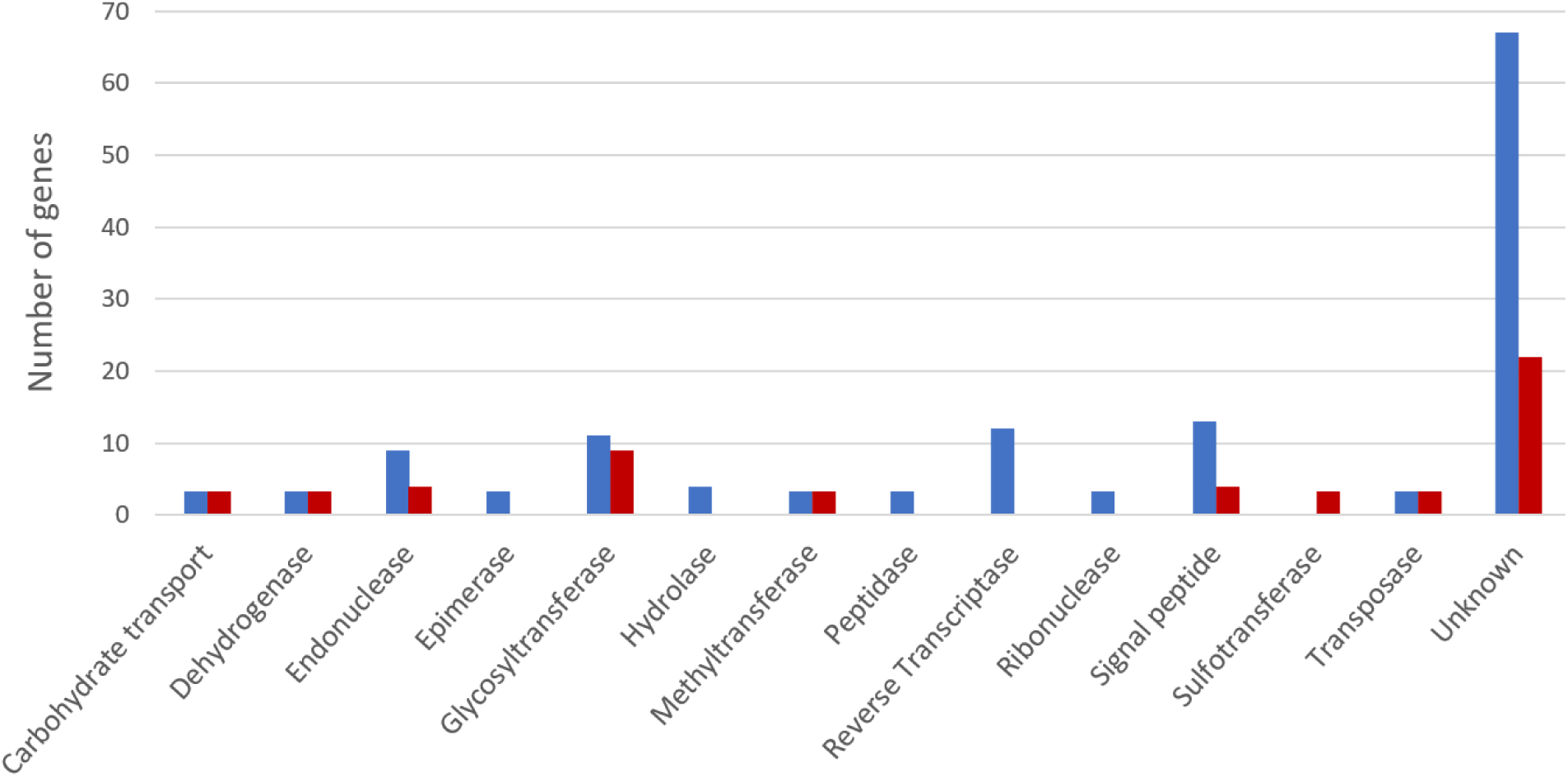
Distribution of biological functions among systematically overexpressed SOC genes across the 4 time points. Blue bars indicate SOC genes overexpressed in the R line, whereas red bars indicate SOC genes overexpressed in the S line.

### Metabolome analyses reveal a shift toward an oxidised lipidome in the susceptible phenotype

The untargeted metabolomic analysis was performed concurrently with transcriptome profiling at *t_1_* and *t_3_* following the protocol described in [25]. A total of 2192 ions were detected. PCA revealed a high variability among individual replicates along the first two principal components, except for the susceptible replicates at *t_1_* (Figure 5A). Despite this inter-replicate variability, the first principal component (PC1, ∼34% of variance) clearly separated samples by the light/dark cycle, with *t_1_* and *t_3_* positioned at opposite ends of the axis (left and right, respectively). The second principal component (PC2, ∼18% of variance) primarily discriminated samples by phenotype: most R replicates were positioned in the lower part of the axis, while most S replicates occupied the upper part. Additionally, the metabolic variability of the S and R lines exhibited an inverse temporal pattern: variability was minimal in the S line at *t_1_* but increased by *t_3_*, while the R line exhibited the inverse pattern. (Figure 5A).

**Figure 5.**
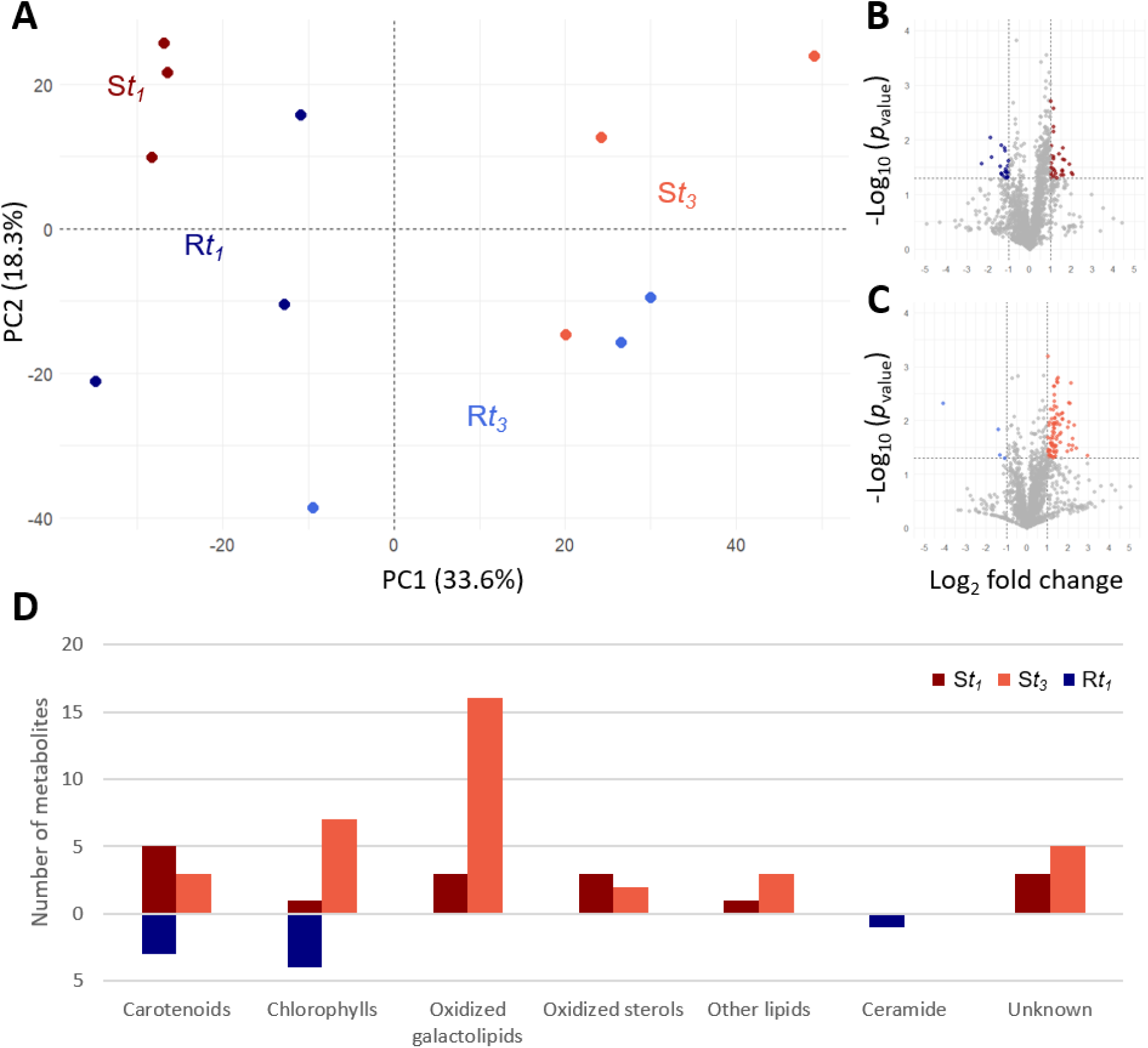
Differential metabolomic analyses and annotation. Comparisons between R (blue) and S (red) lines of *O. mediterraneus* at *t_1_* and *t_3_* respectively. **A**) Principal component analysis of the normalized metabolome of susceptible (S) and resistant (R) *O. mediterraneus* lines at *t_1_* and *t_3_*. Each point represents one sample. **B** and **C**) Volcano plots comparing the 2192 ions between S and R samples. Threshold values: *p*_value_ < 0.05 in the parametric Student’s *t*-test for mean comparison, |log_2_ fold change| ≥ 2. **D**) Histograms of metabolite classes with significantly different abundances between R and S *O. mediterraneus* lines at *t_1_* and *t_3_*.

To identify metabolites contributing to phenotypic variability, we performed a comparative analysis between time points using Student’s *t*-tests on the relative concentrations of detected ions. Ions with a *p*_value_ below 0.05 were considered significantly differentially abundant. To focus on the most biologically relevant changes, we applied a fold change threshold of 2 (|LFC| ≥ 1). The resulting significant ions were then manually curated to remove duplicates (*i.e.*, multiple ions corresponding to the same metabolite) and exclude false positives (Figure 5B and C). Metabolite identification was performed by analysing MS^2^ fragmentation spectra with Sirius 5.6. software [26]. Raw molecular formulas were initially derived from high-resolution MS spectra using FreeStyle 1.6 (Thermo Fisher) and cross-validated with those proposed by Sirius. Candidate natural compounds were identified through manual searches in databases, including the Dictionary of Marine Natural Products (https://dnp.chemnetbase.com) and SciFinder (American Chemical Society). Additionally, MS^2^ spectra were manually inspected and, where necessary, compared against spectral databases, such as the MassBank of North America (MoNA) (https://mona.fiehnlab.ucdavis.edu).

At *t_1_*, this analysis identified 16 metabolites with significantly higher relative concentrations in the S line and 8 metabolites with significantly higher relative concentrations in the R line. At *t_3_*, 36 metabolites exhibited significantly higher relative concentrations in the susceptible line, whereas no metabolites were specifically enriched in the resistant line. Notably, only 4 metabolites were common to both time points, all of which showed higher relative concentrations in the susceptible line. Overall, a total of 56 metabolites differentiated the resistant and susceptible lines across both time points (Table 1). Of these, 48 metabolites were in higher relative concentration in the susceptible line and spanned multiple chemical classes, while the 8 metabolites in higher relative concentration in the resistant line consisted primarily of pigments (Figure 5D).

**Table 1.**
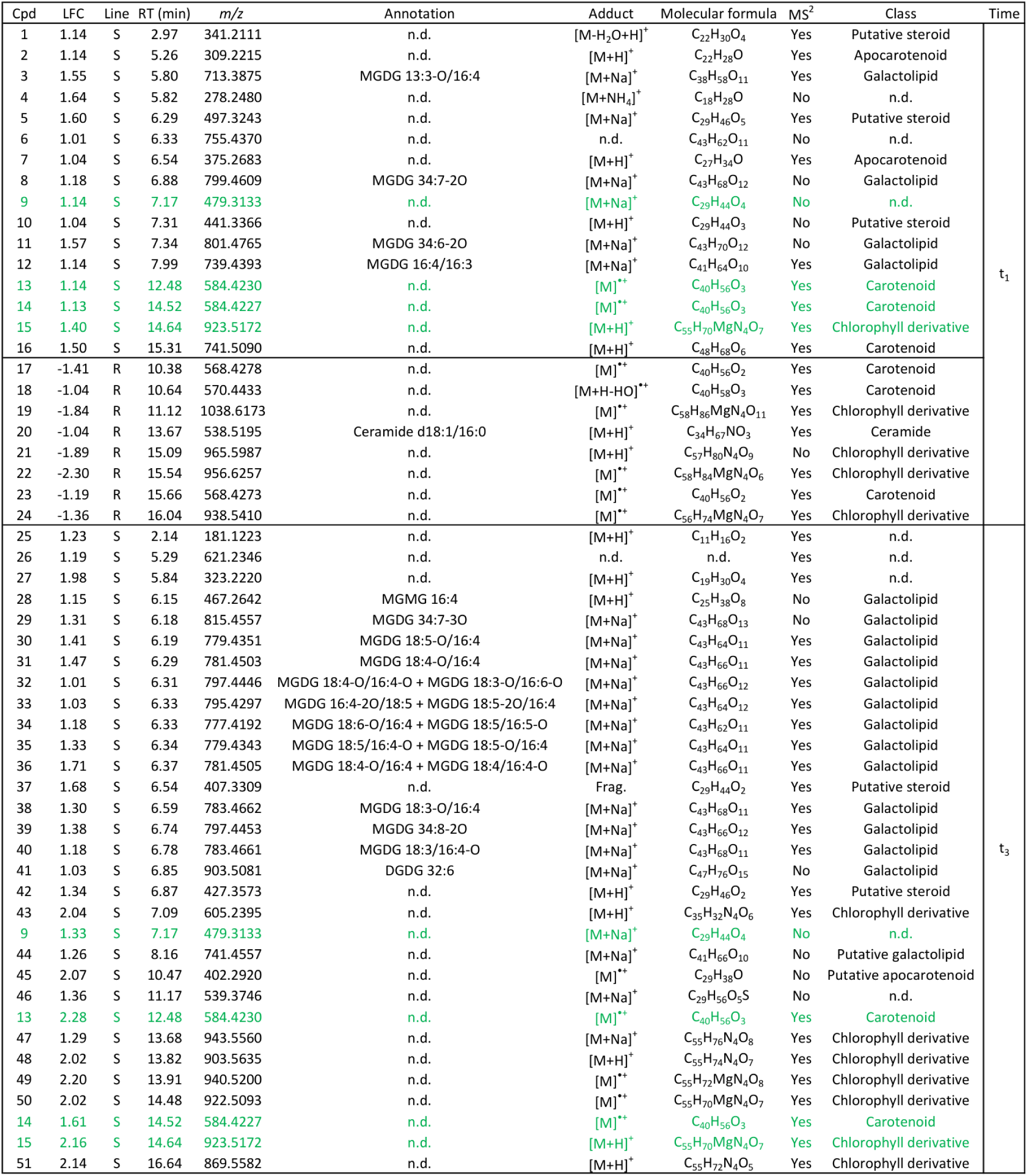
Metabolites whose relative concentration is significantly different between resistant (R) and susceptible (S) lines of *O. mediterraneus*. Common compounds for both times are in green.

At first glance, the metabolome appears less influenced by the S and R phenotype than the transcriptome. Of the 56 differentially abundant metabolites, 49 (87%) were putatively annotated at least at the chemical class level for sterols and pigments. Regiochemical assignments were achieved for most of the lipids, while 7 compounds remained unannotated (unknown).

The metabolomic markers identified belong to five distinct chemical classes, with a predominance of galactolipids, primarily monogalactosyl diacylglycerols (MGDGs). Twenty-three out of the 56 metabolites (41%) were galactolipids, all of which were more abundant in the susceptible line, particularly at *t_3_*. Interestingly, 19 out of the 23 galactolipids were oxidised, with one or two oxidation sites located on the acyl chains at the *sn*-1, *sn*-2, or both positions. The positional assignment (*sn*-1/*sn*-2) of acyl chains on the glycerol backbone of galactolipids was determined by comparing MS^2^ fragmentation patterns. The sodium adduct fragment of MGDGs resulting from the loss of the *sn*-1 acyl chain ([M+Na-R_1_CO_2_H]^+^) exhibited higher intensity than the fragment corresponding to the loss of the *sn*-2 acyl chain ([M+Na-R_2_CO_2_H]^+^) [27]. Based on literature, the galactose moiety of MGDGs was assigned a β(1-3) configuration, while the two galactose units in DGDGs should adopt an α(1-6) conformation [28]. In Table 1, oxidised acyl chains are denoted by an ‘O’ following the carbon number and degree of unsaturation (*e.g.*, 18:4-O for oxidised octadecatetraenoic acid). However, MS data alone do resolve the exact position or type of functional group introduced by oxidation. Four oxidised MGDGs (**8**, **11**, **29** and **39**) were annotated without precise differentiation between their two acyl chains. One oxidised galactolipid, MGDG 13:3-O/16:4 (**3**), contained an uncommon oxidised 13-carbon acyl chain at the *sn*-1 position. This atypical chain length likely results from the activity of a 13-lipoxygenase (LOX-13) followed by a hydroperoxide lyase mediated reaction. Moreover, manual annotation of metabolites **32 to 36**, automatically assigned as one feature, revealed that each of these features were in fact a mixture of two isomers. This was evident from MS^2^ spectra, where two distinct side chain fragmentation patterns were superimposed. For example, both acyl chains of metabolites **32** were oxidised, with different degree of unsaturation of the two galactolipids (MGDG 18:4-O/16:4-O for the **32a** and MGDG 18:3-O/16:6-O for **32b**). In the MS^2^ spectrum, four distinct acyl fragment ions were detected: the highest and lowest *m/z* values matched one possible pair of acyl chains, while the two intermediate fragments formed a second. Both sets were compatible with the precursor ion mass, supporting the presence of two co-eluting isomers. Compounds **33a** and **b** to **36a** and **b** were annotated in the same way. The last four oxidised MGDGs identified (**30**, **31**, **38** and **40**) shared a common structural feature with most MGDGs identified in this study, *i.e.* a C18:n acyl chain at the *sn*-1 position and a C16:4 acyl chain at the *sn*-2 position with oxidations possibly occurring on both acyl chains. The four remaining non-oxidised galactolipids included a digalactosyldiacylglycerol (**41**, DGDG 32:6), a monogalactosyl monoacylglycerol (**28**, MGMG 16:4), a non-oxidised MGDG 16:4/16:3 (**12**), and a putative non-oxidised galactolipid (**44**) annotated only at the class level due to the absence of an MS^2^ fragmentation spectrum. Its identification relied solely on accurate mass and predicted molecular formula. Most galactolipids significantly differentially abundant between susceptible and resistant lines were highly unsaturated and oxidised.

Another important group of marker metabolites comprised oxidised steroids. Five putative steroids were annotated at the class level. Their relative concentrations were significantly higher in the susceptible line compared to the resistant line. These steroids exhibited varying degrees of oxidation, ranging from two (**37** and **42**) to five (**5**) oxygen atoms. The three most oxidised compounds (**1**, **5** and **10)** were detected at *t_1_*, while the remaining two (**37**, **42**) were identified at *t_3_*. Except for compound **1**, which contained 22 carbons, all putative steroids featured a 29-carbon backbone, suggesting derivation from canonical plant sterols such as sitosterol and stigmasterol.

A single ceramide, d18:1/16:0 (**20**), was detected in the analysis and was found to be more abundant in the resistant line at *t_1_*. While this ceramide was the only non-pigment metabolite enriched in the resistant line its isolated occurrence precludes definitive conclusions about its role as a resistance-specific biomarker.

The final two chemical classes identified were pigments, namely carotenoids and chlorophylls. Eleven carotenoids were identified based on their MS^2^ spectra, characterized by the loss of toluene (neutral loss of 92 Da) [29]. In the susceptible line, two apocarotenoids characterised by a shortened polyene chain resulting from oxidative cleavage were annotated at *t_1_* (**2** and **7**), and one putative apocarotenoid at *t_3_* (**45**). Additionally, C_40_ oxygenated derivatives (**13** and **14**) were identified at both time points, alongside a highly oxygenated non-typical putative carotenoid (C_48_H_68_O_6_, **16**). In contrast, the three carotenoids with higher relative concentration in the resistant line were C_40_ carotenoids with fewer oxygens (< 3 oxygens in compounds **17**, **18** and **23**). Twelve chlorophyll derivatives were identified, with one detected in the susceptible line at *t_1_* (**15**), four in the resistant line at *t_1_* (**19, 21, 22** and **24**), and seven in the susceptible line at *t_3_* (**15, 43** and **47 to 51**). Most of these compounds exhibited a characteristic fragmentation pattern, notably the loss of the phytyl chain (C_20_H_38_), resulting in a neutral loss of 278 Da in MS^2^. Within this class, four metabolites (**21**, **47**, **48**, and **51**) were identified as pheophytin derivatives, corresponding to chlorophyll molecules lacking their central metal cation. Furthermore, compound **43** was annotated as a pheophorbide derivative, a chlorophyll derivative missing both the central cation and the phytyl chain. A temporal shift in chlorophyll derivatives distribution was observed: these metabolites were predominantly detected in the resistant line at *t_1_*, whereas they were more abundant in the susceptible line at *t_3_*. Interestingly, at *t_1_*, nearly all identified chlorophyll derivatives contained a central Mg cation, except for compound **21**.

### Integration of transcriptome and metabolome data: how subtle transcriptional shifts may drive profound metabolic shifts

To determine whether differentially abundant metabolites could be directly linked to differentially regulated genes, we examined the top 1,000 genes contributing most significantly to principal component 4 of the PCA, which distinguishes the resistant (R) and susceptible (S) phenotypes (Figure 1B). Among these genes, only one gene (Ostme09g02170) was directly associated with a biomarker biosynthetic pathway: it encodes phosphatidic acid phosphatase (PAP), a key enzyme in the oxidized galactolipid pathway (Figure 6).

**Figure 6.**
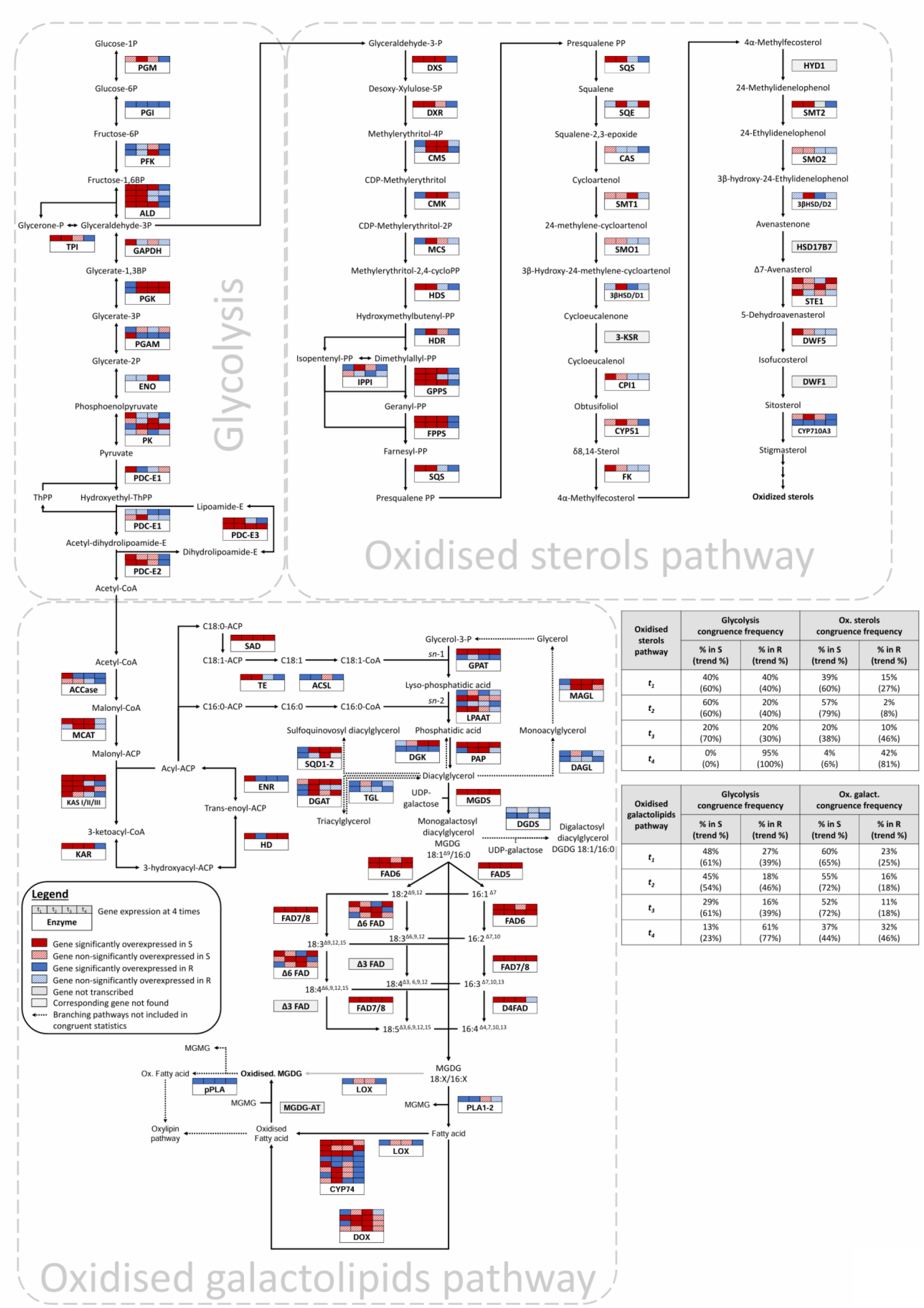
Temporal expression dynamics and phenotype-associated congruence across glycolysis, oxidised galactolipid, and oxidised phytosterol biosynthetic pathways, with expression profiles of all enzymatic steps, including: phosphoglucomutase (PGM), phosphoglucose isomerase (PGI), phosphofructokinase (PFK), aldolase (ALD), triose phosphate isomerase (TPI), glyceraldehyde-3-phosphate dehydrogenase (GAPDH), phosphoglycerate kinase (PGK), phosphoglycerate mutase (PGAM), enolase (ENO), pyruvate kinase (PK), pyruvate dehydrogenase complex subunits (PDC-E1, PDC-E2, PDC-E3), and thiamine pyrophosphate (ThPP); acetyl-CoA carboxylase (ACCase), malonyl-CoA-acyl carrier protein transacylase (MCAT), 3-ketoacyl-ACP synthases I/II/III (KASI/II/III), ketoacyl-ACP reductase (KAR), 3-hydroxyacyl-ACP dehydratase (HD), enoyl-ACP reductase (ENR), stearoyl-ACP desaturase (SAD), thioesterase (TE), long-chain acyl-CoA synthetase (ACSL), glycerol-3-phosphate acyltransferase (GPAT), lysophosphatidic acid acyltransferase (LPAAT), phosphatidic acid phosphatase (PAP), diacylglycerol kinase (DGK), diacylglycerol lipase (DAGL), monoacylglycerol lipase (MAGL), sulfoquinovosyl diacylglycerol synthase 1 and 2 (SQD1-2), diacylglycerol acyltransferase (DGAT), triacylglycerol lipase (TGL), monogalactosyldiacylglycerol synthase (MGDS), digalactosyldiacylglycerol synthase (DGDS), and fatty acid desaturases (FAD); phospholipase A1/A2 (PLA1-2); lipoxygenase (LOX); cytochrome P450, family 74 (CYP74); dioxygenase (DOX); monogalactosyldiacylglycerol acyltransferase (MGDG-AT); patatin-like phospholipase (pPLA); 1-deoxy-D-xylulose-5-phosphate synthase (DXS), 1-deoxy-D-xylulose 5-phosphate reductoisomerase (DXR), 4-diphosphocytidyl-2-C-methyl-D-erythritol synthase (CMS), 4-diphosphocytidyl-2-C-methyl-D-erythritol kinase (CMK), 2-C-methyl-D-erythritol 2,4-cyclodiphosphate synthase (MCS), 1-hydroxy-2-methyl-2-(E)-butenyl-4-diphosphate synthase (HDS), 4-hydroxy-3-methylbut-2-enyl diphosphate reductase (HDR), isopentenyl diphosphate isomerase (IPPI), geranyl diphosphate synthase (GPPS), farnesyl diphosphate synthase (FPPS), squalene synthase (SQS), squalene epoxidase (SQE), cycloartenol synthase (CAS), sterol methyltransferases 1 and 2 (SMT1, SMT2), sterol methyl oxidases 1 and 2 (SMO1, SMO2), 3β-hydroxysteroid dehydrogenases/decarboxylases 1 and 2 (3βHSD/D1, 3βHSD/D2), 3-ketosteroid reductase (3-KSR), cyclopropyl isomerase (CPI1), sterol C14-demethylase (cytochrome P450, family 51 or CYP51), Δ14-sterol reductase (FK), C8-C7 sterol isomerase (HYD1), sterol C-7 reductase (STE1), Δ7-sterol-C5-desaturase (DWF5), Δ24-sterol reductase (DWF1), and sterol C22 desaturase (cytochrome P450, family 710-A3 or CYP710A3).

We hypothesized that the lack of overlap between the differentially expressed genes and differentially abundant metabolites may be the consequence of the involvement of DEGs in gene expression regulation (*e.g.*, transcription factors, TEs), rather than in biosynthetic pathways, as such pathways are tightly regulated to maintain cellular homeostasis. To explore this further, we investigated whether more subtle pathway-level transcriptional changes could explain the observed differences in biomarker accumulation.

Whole genome data and annotation enabled the identification of the candidate genes involved in most reactions of the well-characterized metabolic pathways upstream of several biomarker production pathways (Figure 6). While kinetic information for individual reactions within these pathways remains unknown, we applied a heuristic approach to compare the transcriptomic and metabolomic profiles of the R and S phenotypes. Briefly, we hypothesized that increased biomarker levels could arise from either (i) higher efficiency in upstream reactions leading to the biomarker synthesis, or (ii) lower efficiency in downstream reactions processing the biomarker. The higher efficiency upstream reactions might be due to either an increased transcription rate of one of the enzymes involved, and/or reduced negative feedback on any of these enzymes. The decrease in downstream reactions processing the biomarker may be due to a decreased transcription rate of one of the enzymes involved, and/or reduced positive feedback against any of these enzymes. Given the lack of information on feedback loops, we focused on relative changes in transcription rates of the enzyme involved directly in the pathway. We considered all differentially expressed genes with significant changes (*p*_adj_ < 0.05), regardless of fold-change magnitude, even those below our initial thresholds (|log_2_ fold change| ≥ 2). To evaluate pathway-level congruence, we calculated the frequency of consistent transcriptional changes between the R and S phenotypes. For biomarkers overrepresented in S, this was determined by estimating the proportion of genes with significantly higher transcription rates in S relative to the total number of candidate genes in each pathway (Figure 6).

We focused on the glycolysis, galactolipid and phytosterol pathways, as these upstream metabolic pathways lead to the biosynthesis of the oxidised galactolipid and sterols, which are biomarkers of the virus-susceptible phenotype.

#### Upstream glycolysis pathway shows moderate congruence

Glycolysis provides central metabolic precursors that feed into multiple biosynthetic pathways, including sterol and galactolipid synthesis, from which several oxidised derivatives were identified as biomarkers enriched in the S phenotype. To explore potential transcriptional congruence between glycolysis and the biosynthesis of these biomarkers, we analysed the expression patterns of glycolytic genes across all four time points.

From *t_1_* to *t_3_*, we observed a moderate transcriptional bias towards overexpression in the S phenotype. Specifically, the proportion of glycolytic genes significantly overexpressed in S was consistently higher than in R with 48%, 45% and 29% overexpressed in S compared to 27%, 18% and 16% in R at *t_1_*, *t_2_* and *t_3_*, respectively (Figure 6). This trend was further supported when including non-significant trends, with 61%, 54%, and 61% of genes were overexpressed in S at *t_1_*, *t_2_*, and *t_3_* respectively, compared to 39%, 46%, and 39% in R. These patterns suggest a subtle but consistent predominance in gene expression supporting the S-associated metabolic phenotype during the first three timepoints of the experiment. However, this congruence vanished at *t_4_*, where a sharp transcriptional shift occurred: 61% of glycolytic genes were significantly overexpressed in the R phenotype, with 77% genes trending in the same direction.

This analysis included key enzymes catalysing rate-limiting steps in glycolysis, such as phosphofructokinase (PFK) and pyruvate kinase (PK), which provide key insights into the regulation of carbon flux toward downstream biosynthetic pathways. Notably, the expression of PFK and PK genes varied over time, but both enzymes ultimately showed a stronger overexpression in the R phenotype, particularly at *t_4_*.

#### Oxidised galactolipids pathway

Similarly, the galactolipid biosynthesis pathway, which relies on glycolysis-derived precursors such as acetyl-CoA, G3P or UDP-Galactose, showed signs of transcriptional alignment with the metabolomic profiles characteristic of the S phenotype. This pathway is responsible for the synthesis of MGDG, a major thylakoid lipid from which several oxidised forms were enriched in susceptible cells (Figure 6).

Out of the 30 genes involved in the synthesis of these galactolipids, homologous candidate genes could be identified except for gene Δ3 FAD and MGDG-AT (Figure 6, Table S10). When they were several candidates for the same genes, all were included (Figure 6). For example, two putative DGDS-encoding genes were identified, both of which exhibited either significant overexpression or a consistent trend toward the R phenotype across all time points. This is consistent with a metabolic flux preferentially directed toward MGDG synthesis in the S phenotype. In *Ostreococcus*, newly synthesised galactolipids such as MGDG 18:0/16:0 and MGDG 18:1/16:0 are relatively minor components, while highly unsaturated MGDGs predominate. Their formation requires sequential desaturation of both acyl chains, catalysed by fatty acid desaturases (FADs). Six FADs have been previously characterised in the *Ostreococcus* genus: a stearoyl-ACP desaturase (SAD), an acyl-CoA Δ6 desaturase localised to the endoplasmic reticulum (ER) in *O. tauri* [30], a Δ4 FAD from the ER of *O. lucimarinus* [31], a Δ5 FAD from the ER of both *O. lucimarinus* and *O. tauri* [32], and two recently identified plastidial desaturases of the Δ6-ω6 and Δ6-ω3 types [33]. The majority of these desaturases displayed expression patterns associated with the S phenotype over time, consistent with enhanced galactolipid unsaturation.

From *t_1_* to *t_3_*, the transcriptional profiles of genes involved in galactolipid biosynthesis revealed a strong and consistent skew toward overexpression in the S phenotype. The proportion of significantly overexpressed genes in S was 60%, 55%, and 52% at *t_1_*, *t_2_*, and *t_3_*, respectively, while the corresponding values for R were substantially lower at 23%, 16%, and 11%. This pattern was further supported by non-significant expression trends, where 65%, 72%, and 72% of genes exhibited a trend toward higher expression in S at the same time points. Collectively, these results reflect a coordinated transcriptional upregulation of the galactolipid biosynthetic pathway in susceptible cells. At *t_4_*, this transcriptional bias diminished markedly. Only 37% of genes remained significantly overexpressed in S, while the proportion of genes significantly overexpressed in R rose to 32%.

Several key enzymatic steps in this pathway, like acetyl-CoA carboxylase (ACCase), glycerol-3-phosphate acyltransferase (GPAT), phosphatidic acid phosphatase (PAP), fatty acid desaturase (FAD) or MGDG synthase (MGDS), are known to exert regulatory control over metabolic flux. These enzymes exhibited distinct transcriptional dynamics over time. ACCase genes showed a predominant overexpression in the R phenotype, particularly from *t_2_* to *t_4_*, while earlier time points displayed either non-significant trends or S-biased gene expression. In contrast, MGDS, PAP and most FAD genes were uniformly overexpressed in the S phenotype from *t_1_* to *t_4_*, mirroring the accumulation of oxidised galactolipids in susceptible cells. GPAT expression appeared more heterogeneous, with one candidate gene consistently overexpressed in the S phenotype and another candidate being overexpressed in the R phenotype. Altogether, these transcriptional profiles indicate a tightly regulated and temporally restricted activation of galactolipid biosynthesis in S cells, which is progressively lost in favor of a resistance-associated state at the end of the experiment.

The oxidation of galactolipid acyl chains, leading to the formation of oxidised MGDGs, has not been elucidated in *O. mediterraneus*. One hypothesis involves the direct oxidation of galactolipids by lipoxygenases. In support of this, a single putative LOX gene (Ostme02g00890) was identified based on the presence of a PLAT/LH2 domain characteristic of this enzyme class. However, its expression was variable across time points, with significant overexpression in the R phenotype at *t_1_* and *t_4_*, and a trend towards higher expression in S at *t_2_* and *t_3_*. The comparative transcription rates in R and S of this candidate gene thus poorly explains the accumulation of oxidised galactolipids in the S lines. An alternative hypothesis involves the hydrolysis of acyl chains, followed by their oxidation and re-esterification onto glycerol backbones. Similarly, one homolog of phospholipase (PLA1-2, Ostme05g01300) was identified in *O. mediterraneus* and showed consistent overexpression in the R phenotype at *t_1_*, *t_2_* with a trend at *t_4_.* However, eight putative oxygenases with CYP74-related domains were found, including two that were significantly overexpressed in the S phenotype from *t_1_* to *t_3_*, and one that was overexpressed at *t_1_* and *t_4_* with a trend toward S at *t_2_* and *t_3_*. In addition, among four annotated α-dioxygenase (α-DOX) genes, most were either significantly overexpressed or with an overexpression trend toward the S phenotype, suggesting that oxidase activity may be enhanced in susceptible cells, potentially accounting for the increased accumulation of oxidised MGDGs. Interestingly, the R phenotype significantly overexpressed an orthologous gene to pPLA in *A. thaliana* (Ostme11g03410) throughout the time series. This pPLA gene encodes a patatin-like phospholipase implicated in the hydrolysis of oxidised acyl chains from galactolipids and the release of oxylipin precursors such as (dn)OPDA for jasmonate biosynthesis. Its overexpression in the R phenotype could thus contribute to the reduced levels of oxidised galactolipids in the R phenotype, possibly through active degradation or remodelling of damaged membrane lipids, while the lower pPLA activity in S may favour their accumulation.

Oxidised fatty acids are expected to feed into the oxylipin pathway, which partly overlaps with β-oxidation. Core β-oxidation enzymes implicated in oxylipin catabolism were identified (Table S11), including two candidate acyl-CoA oxidases (ACX, Ostme01g04280), predominantly overexpressed in R (*t_1_*, *t_2_*, and *t_4_*), and Ostme07g02190, more strongly induced in S (*t_2_* and *t_4_*); Ostme07g03180 a single multifunctional protein (MFP2), consistently overexpressed in R across all time points; and Ostme05g04530, a thiolase (KAT) upregulated in R at *t_1_* and in S at *t_3_* and *t_4_*. At t1, the overexpression of the 3 genes in R support a higher rate of synthesis of oxylipin precursors in R, consistent with our observation that oxidized fatty acids are biomarkers of the S phenotype.

Beyond the oxylipin pathway, the resistant line also activated other genes involved in lipid turnover, storage, and remodelling (Table S10). Notably, diacylglycerol lipase (DAGL) and triacylglycerol lipase (TGL) genes were consistently overexpressed, suggesting active lipid catabolism (Figure 6). Monoglyceride lipase (MAGL) genes exhibited an early peak in the resistant line (*t_1_*), followed by a shift toward the susceptible phenotype at later time points. Diacylglycerol diacylglyceryltransferase (DGAT) genes and diacylglycerol glycosyltransferase (DGDS) genes showed mixed but predominantly over-expression in the R lines. In addition, enzymes involved in sulfoquinovosyl diacylglycerol (SQDG) synthesis, namely SQD1 and SQD2, were upregulated in the resistant line at early stages (*t_1_*-*t_2_*), preceding a transcriptional shift toward the susceptible phenotype (Figure 6).

#### Phytosterols pathway

In parallel, we examined the transcriptional congruence of the oxidised sterol biosynthesis pathway, which also depends on glycolysis-derived precursors. This pathway leads to the synthesis of stigmasterol and then oxidised sterols identified as a biomarker in the S phenotype. Most genes (27 out of 30) involved in this pathway could be identified (Table S10).

The expression dynamics of the oxidised phytosterol biosynthetic pathway revealed a strong and temporally structured transcriptional overexpression in the S phenotype during the early daytime sampling points. At *t_1_* and *t_2_*, the proportion of genes significantly overexpressed in S was markedly higher than in R, reaching 39% and 57% for S compared to only 15% and 2% for R, respectively. These differences were reinforced by non-significant trends, with 60% and 79% of genes trending toward overexpression in S, against 27% and 8% in R. At *t_3_*, however, the pattern became more ambiguous: while the proportion of significantly overexpressed genes remained higher in S 20% than in R 10%, the trend shifted slightly in favour of R with 46%, suggesting a transitional transcriptional state. This partial reorientation became more pronounced at *t_4_*, where congruence of the metabolomic and the transcriptomic profiles in the S phenotype collapsed entirely: only 4% of genes were significantly overexpressed in S, compared to 42% in R. Altogether, these results indicate a temporally constrained transcriptional activation of the phytosterol biosynthetic pathway in S cells, which is progressively reversed in favour of a resistance-associated profile during the late phase of the experiment.

Key regulatory bottlenecks within the phytosterol biosynthesis pathway include enzymes of the methylerythritol phosphate (MEP) pathway, which provides plastid-derived isoprenoid precursors. Among them, 1-deoxy-D-xylulose-5-phosphate synthase (DXS), 1-deoxy-D-xylulose-5-phosphate reductoisomerase (DXR), 4-hydroxy-3-methylbut-2-enyl diphosphate synthase (HDS), and 4-hydroxy-3-methylbut-2-enyl diphosphate reductase (HDR) are widely recognised as rate-limiting steps and major control points of carbon flux into sterol biosynthesis [34,35]. Downstream, additional checkpoints within the sterol-specific segment are represented by squalene synthase (SQS), squalene epoxidase (SQE), and cycloartenol synthase (CAS), together with branching enzymes such as sterol methyltransferases (SMT1 and SMT2) and C-22 sterol desaturase (CYP710A3), which determine the formation of phytosterol end-products including sitosterol and stigmasterol. The expression profiles of these genes largely mirrored the global transcriptional trends of the pathway, with a strong predominance of overexpression in S at early time points (*t_1_*-*t_2_*), a more mixed profile at *t_3_*, and a consistent overexpression in R at *t_4_*. These temporal patterns suggest that transcriptional regulation of flux-controlling enzymes may underlie the early activation of sterol oxidation in susceptible cells and its progressive suppression in favour of a resistance-associated metabolic state.

#### The search of sterol oxidases

The oxidised sterol biomarkers identified in this study exhibit between two and five oxidation events, suggesting the involvement of multiple enzymatic steps, likely catalysed by sterol oxidases. However, the conserved sequence motifs typically associated with sterol oxidase activity, are also found in related enzyme classes, including sterol desaturases and fatty acid hydroxylases. Consequently, although six candidate genes were identified based on the presence of the InterPro domain IPR006694, characteristic of the fatty acid hydroxylase family that also encompasses C-5 sterol desaturases and C-4 sterol methyl oxidases (SMOs), their precise biochemical function remains unresolved (Table S12). Three out of these six candidate genes exhibited strong and significant overexpression in the susceptible phenotype at early time points (*t_1_* and/or *t_2_*), mirroring the temporal accumulation of oxidised sterol biomarkers in this condition. In contrast, one candidate displayed consistent and significant overexpression in the resistant phenotype at later time points (*t_2_* to *t_4_*), while two candidates showed either non-significant or inconsistent expression patterns. We thus hypothesize that the most likely candidate genes involved in phytosterol oxidation in susceptible cells are Ostme11g00750, Ostme03g01060, and Ostme12g01770. However, alternative enzymatic or non-enzymatic mechanisms, not yet identified, may also contribute to the biosynthesis of these highly oxidised sterol derivatives.

Overall, while glycolysis serves as the central upstream provider of carbon skeletons for both sterol and galactolipid biosynthesis, its transcriptional regulation exhibits only moderate phenotype-specific congruence. In contrast, downstream biosynthetic pathways, particularly those leading to oxidised sterols and galactolipids, display a much stronger and temporally structured transcriptional alignment with the accumulation of biomarkers in the S phenotype. This transcriptomic-metabolomic congruence diminishes at night, with genes involved in sterol biosynthesis more expressed in the R phenotype. Collectively, these results highlight phenotype-specific metabolic reprogramming, most pronounced in sterol metabolism. While upstream glycolytic flux is broadly modulated, the selective transcriptional activation of rate-limiting enzymes in downstream pathways appears to play a decisive role in shaping phenotype-specific metabolic outcomes.

## Discussion

### Transcriptional changes on the small outlier chromosome in resistant and susceptible lines

The small outlier chromosome (SOC) is a unique, hypervariable chromosome described in all sequenced Mamiellales genomes, including *Ostreococcus*, *Micromonas*, and *Bathycoccus* [14,36,37]. In *Ostreococcus*, it exhibits extreme structural and transcriptional plasticity [13,38] – particularly in response to viral infections [14]. A previous comparative transcriptomic analysis in virus-resistant *Ostreococcus tauri* lines revealed a bipartite gene expression structure on the SOC [14] with a 100 kb region expressed in the S lines, and a 180 kb region expressed in R lines. Here, we extend this finding to *O. mediterraneus*, where the SOC (∼640 kb) is more than twice the size of that in *O. tauri* (∼290 kb) [11]. Our transcriptional analyses confirm the presence of phenotype-specific expression patterns on the SOC, with a ∼110 kb region associated with the S phenotype and a ∼530 kb region linked to the R phenotype. Notably, our experimental setup provides evidence that the SOC transcriptional changes remain stable across the diel cycle in *O. mediterraneus*, while it is not the case for most DEGs on the standard chromosomes.

Consistent with previous observations in other species [4], there are only seven orthologous SOC genes between *O. mediterraneus* and *O. tauri*, none of which have functional annotations. Five of these genes rank among the top 300 contributors to the separation of resistant and susceptible phenotypes along PC4 of the PCA (Figure 1B). Despite this lack of sequence conservation, the SOC consistently harbours a few genes with similar predicted functions, such as glycosyltransferases (GTs) and methyltransferases [13,14,37]. This supports a scenario of functional convergence of the genes recruited on the SOC [37]. Strikingly, there was a higher expression of genes with predicted sulfotransferase activity in the S lines of *O. tauri* and *O. mediterraneus*. Sulfotransferases catalyse the transfer of sulfuryl groups (SO_3_^-^) to diverse acceptors, including hormones, metabolites, and membrane components thereby modulating their activity, stability, and cellular fate [39]. However, all 20 predicted GTs are differentially expressed both in R and S lines in *O. mediterraneus*, suggesting extensive remodelling of glycoconjugates. Glycosylation in eukaryotes occurs predominantly in the ER and the Golgi apparatus, which together constitute central hubs for the synthesis and remodelling of polysaccharides and for *N*-glycosylation, a tightly regulated process essential for protein folding, quality control and plant immune responses [40–43]. For example, in *Arabidopsis thaliana*, deletion of the ER-localised α-1,3-mannosyltransferase gene ALG3 disrupts the glycosylation of the secreted effector protein Slp1 (Secreted LysM Protein1), preventing immune suppression by *Magnaporthe oryzae* and maintaining host resistance [44]. Four GTs overexpressed in the S line are physically clustered on the SOC, suggesting co-regulation and functional convergence within a shared biosynthetic pathway. Their membrane-associated localisation and shared functional domains support a possible role in the biosynthesis or remodelling of extracellular glycoconjugates, such as glycosaminoglycans. Since glycosaminoglycans have been implicated as viral attachment factors in other biological systems [45–47], we hypothesise that the overexpression of these genes in the S line might facilitate viral entry. Surface glycans play a key role in host-virus interactions by mediating the initial steps of binding and recognition between host cells and viral particles at the cell surface [48]. An alternative hypothesis could be that the overexpression of specific SOC GT confers resistance by competing with viral GTs for shared glycosylation substrates, thereby disrupting viral assembly. Previous studies in another dsDNA virus, Chlorella virus PBC, demonstrated that the capsid of the V-1 was heavily glycosylated [49]. Given that prasinovirus genomes frequently encode their own GTs—some of which are conserved across species—it is plausible that a similar glycosylation-dependent process occurs in prasinoviruses, further emphasizing the critical role of glycosylation in viral infection. Further glycobiological studies will be essential to elucidate the role of SOC-encoded GTs in viral susceptibility and resistance.

### Diel and cell cycle transcriptional shifts underlie R/S phenotype differences

In sharp contrast to the stable R/S gene expression changes for genes on the SOC chromosome; there are only 6 out of 213 DEGs on the standard chromosomes that are systematically differentially expressed across all 4 time points. In the S line, early time point (*t_1_*) overexpression of genes involved in nuclear processes, including transcription and RNA metabolism, suggests an increased cellular activity potentially linked to growth initiation. A transient enrichment of geranylgeranyl reductase expression was observed in the susceptible line at *t_3_*, a timepoint corresponding to the early night phase. Geranylgeranyl reductase mediates the production of phytyl-PP, an essential precursor for chlorophylls and tocopherols [50]. This temporal pattern may reflect anticipatory chlorophyll turnover or enhanced isoprenoid metabolism in preparation for the following photoperiod [51].

In the R line, there is an enrichment of genes related to nitrogen metabolism at *t_3_*, indicating a possible enhanced capacity for nitrogen assimilation or processing that could support metabolic robustness. Interestingly, this transcriptional activation occurs at the beginning of the night, a period during which nitrogen metabolism is typically limited by the absence of light in *Ostreococcus* [22]. During viral infection of *O. tauri*, host nitrogen metabolism genes are strongly repressed at night while viral transcription increases sharply [52]. The fact that the R line maintains or even enhances nitrogen assimilation-related gene expression during this phase, despite the absence of viral stress and under nitrogen-rich conditions, suggests a decoupling from diel repression patterns. This atypical activation may underlie a protective role, as nitrate reduction is known to generate nitric oxide (NO), a reactive signaling molecule involved in triggering cellular defense responses. Both NO and ROS (reactive oxygen species), which are tightly connected through redox signaling, are known to mediate antiviral responses in higher plants such as *Arabidopsis* [53], and in microalgae like *Gephyrocapsa huxleyi* [54]. Their production through nitrogen metabolic activity may thus represent a preemptive strategy in the R line to maintain a primed defensive state, enhancing its capacity to withstand future viral encounters. By the final timepoint (*t_4_*), resistant cultures show pronounced enrichment of ion transport genes, especially those involved in potassium ion homeostasis, likely contributing to the maintenance of cellular osmotic balance and membrane potential, which are critical for cellular stability [55]. Such robust cellular homeostasis, particularly through efficient ion regulation, is a key strategy for mitigating cellular stress and thereby limiting the production of deleterious ROS [56]. Several genes located on the standard chromosomes, particularly those involved in nitrogen metabolism, redox regulation, and ion transport, emerge as putative biomarkers of viral resistance. While these genes likely reflect constitutive physiological changes, their temporal dynamics and functional relevance suggest a potential role in preemptive stress buffering or immune priming. A more comprehensive picture emerges when transcriptomic signals are considered alongside metabolomic data, especially within central carbon and lipid metabolic pathways.

Integrated transcriptomic and metabolomic analyses reveal phenotype-specific metabolic reprogramming, with diel cycles and cell cycle progression shaping lipid metabolism and transcription-metabolite coupling. In the susceptible (S) phenotype, transcriptional activation of lipid pathways peaks during the daytime (*t_1_*, *t_2_*), aligning with the light phase. MGDG biosynthesis in the S line exhibits strong transcription-metabolite congruence at *t_1_* and *t_3_*, but the high oxidised galactolipid levels at *t_3_*—despite transcriptional downregulation—may indicate a temporal lag between gene expression and metabolite turnover. These temporal lags and decoupling align with cell cycle-dependent metabolic fluxes [57,58] and highlight phenotype-specific metabolic strategies, possibly modulated by light-dark transitions. Such phase shifts in metabolic activity may influence virus susceptibility, as viral replication in *Ostreococcus* is timing-dependent: infections during the light phase trigger rapid lysis, while late-day infections delay lysis by ≥ 48 hours [52]. This suggests that host metabolic rhythms may critically influence viral success, with phase-shifted metabolism of the R line potentially serving as a preemptive antiviral mechanism.

### Oxidised Galactolipids in virus susceptible *Ostreococcus mediterraneus*

This study provides the first characterization of oxidised monogalactosyldiacylglycerol (MGDG) species in this marine picoeukaryote. Several oxidised MGDG species were differentially abundant across lines and time points, yet they were biomarkers of the S line, indicating a line-specific regulation of galactolipid oxidation in *O. mediterraneus*. Notably, these oxidised forms correspond to the major non-oxidised MGDG species previously described in this alga, namely MGDG 18:3/16:4, 18:4/16:4, and 18:5/16:4 [25,59]. In higher plants species such as *A. thaliana*, oxidised galactolipids, particularly oxidised forms of MGDG, have been investigated for their roles in signalling and defence responses [60–63]. Arabidopsides are the most well-characterised oxidized galactolipids and are thought to act as precursors or storage forms for jasmonate-family signals while free oxidised fatty acids released by lipases can act as signalling molecules to modulate transcription responses [63–65]. Interestingly, certain oxidised galactolipids display enhanced bioactivity compared to their free oxylipin counterparts [61], suggesting a functional role for the glycolipid scaffold itself.

Here, transcriptional and metabolic integration reveals that the biosynthesis of oxidised MGDGs in *O. mediterraneus* is different between R and S lines. The S line exhibited strong transcriptional activation of monogalactosyldiacylglycerol synthase (MGDS) and fatty acid desaturases (FAD5, FAD7, FAD8), enhancing flux toward MGDG production and subsequent oxidation. This coordinated transcriptional activation was tightly coupled with metabolite accumulation, as reflected by the pronounced enrichment of oxidised MGDGs at *t_1_* and *t_3_*. In parallel, the R line displayed upregulation of acetyl-CoA carboxylase (ACCase), a recognised regulatory bottleneck in plastidial fatty acid biosynthesis, particularly from *t_2_* to *t_4_*, suggesting an enhanced entry of carbon flux into the lipid pathway. The R line also exhibited early upregulation of patatin-like phospholipase (pPLA), suggesting active degradation of oxidised galactolipids. Although a canonical oxylipin pathway appears incomplete in *O. mediterraneus*, with no clear orthologues of the first biosynthetic enzymes, the core β-oxidation machinery implicated in oxylipin catabolism was detected, indicating that the cycle is at least partially functional. Notably, candidate genes encoding ACX, MFP2, and KAT were preferentially expressed in the R line, particularly at early time points, consistent with a more active turnover of oxidised fatty acids. While it remains uncertain whether *O. mediterraneus* is capable of synthesising oxylipins such as 12-oxo-phytodienoic acid (OPDA) or jasmonic acid (JA), the well-established signalling functions of these molecules in vascular plants provide a valuable comparative framework. For instance, OPDA and JA are well-established mediators of plant defence, with JA functioning as a central immune hormone [66] and OPDA emerging as an independent regulator capable of modulating gene expression without conversion to JA-Ile [67]. OPDA contributes to protective responses against both fungal pathogens and insect herbivores [68,69], in part through its conjugation with isoleucine to form OPDA-Ile, a bioactive compound with transcriptional activity [70]. The idea that oxidised acyl chains or MGDGs could function analogously to oxylipins in plant defense is supported by recent studies in grapevine, where resistant phenotypes to *Plasmopara viticola* displayed a more oxidised lipidome and a distinct activation of plastidial phospholipase A during infection [71,72]. Additional transcriptional activation of lipases (TGL, DAGL) and alternative lipid pathways (DGDS metabolism) further underscore lipid turnover and membrane remodelling as central features of the resistant phenotype in *Ostreococcus*. Taken together, these findings, in line with recent studies in other lineages, suggest that oxidised MGDGs may not only accumulate passively but could also function as regulatory metabolites in stress and defence, paralleling the role of oxylipins in vascular plants. In this framework, the R line of *O. mediterraneus* may actively remodel and catabolise oxidised galactolipids to repurpose them for protective or signalling functions, while the S line appears limited to passive accumulation, which could underlie its differential susceptibility to viral infection.

### Putative Sterol Involvement in Host-Virus Interactions in *O. mediterraneus*

In addition to MGDG, oxidised sterol compounds were also found to be more abundant in the S line. Sterols are essential components of eukaryotic membranes, where they regulate fluidity, permeability, and interactions with membrane proteins and lipids [73]. Beyond their structural roles, they act as precursors for a wide range of bioactive molecules involved in cellular and developmental processes [74] while in plants they have also been associated with cell proliferation, signal transduction, and the regulation of membrane-bound enzyme activities [73]. Sterols further participate in more specific cellular processes, including gene expression regulation and host-virus recognition [75]. The role of sterols in algal-virus interactions was first demonstrated in the *Gephyrocapsa huxleyi*-GhV model, where lipidomics revealed host sterol depletion during infection and incorporation of specific sterols into the viral membrane [15]. Pharmacological inhibition of the host mevalonate pathway confirmed that sterols are hijacked for virion assembly. Such exploitation appears conserved, as West Nile Virus upregulates cholesterol biosynthesis in human cells [76], while hosts can counteract by repressing sterol synthesis, as observed in interferon-mediated restriction of cytomegalovirus replication [77]. Together, these observations highlight sterols as central players in host-virus interactions, either as structural resources co-opted for virion assembly or as regulatory metabolites modulated by the host to restrict infection. In *O. mediterraneus*, the differential abundance of sterol-like compounds between R and S lines suggests that variations in sterol metabolism could influence viral success. Future characterisation of the sterol composition of virus particles will therefore be essential to determine whether sterols act as passive structural elements or as active determinants of susceptibility and resistance.

The oxidised phytosterol biosynthetic pathway exhibited a strong and temporally structured transcriptional expression between phenotypes. The susceptible line displayed daytime overexpression, consistent with an initial orientation toward sterol biosynthesis and oxidation, whereas the resistant line exhibited delayed but sustained activation at the end of night, suggesting temporally segregated engagement of sterol metabolism. The expression of the four oxysterol-binding proteins (OSBPs) identified in *Ostreococcus* (Table S13), including several species-specific paralogs, displayed phenotype-dependent temporal patterns, suggesting distinct dynamics of sterol transport and membrane remodelling in *O. mediterraneus*. Among them, the ortholog of *A. thaliana* ORP3A (Ostme09g00040), implicated in sterol trafficking to the plasma membrane [78], showed an early induction in the S line followed by sustained expression in the R line, pointing to temporally segregated regulation of sterol fluxes. Another OSBP-related gene (Ostme16g02140), lacking homologs in other Mamiellophyceae, was strongly expressed in resistant lines and may represent a lineage-specific innovation linked to antiviral defense in *O. mediterraneus*. OSBP and OSBP-related proteins (ORPs) form a highly conserved family across eukaryotes [79]. Given that sitosterol and stigmasterol associate with glycosyl inositol phosphoceramides (GIPCs) to form lipid rafts, dynamic membrane microdomains enriched in signaling proteins and enzymes [80,81], transcriptional shifts in sterol metabolism are expected to reshape raft composition and receptor accessibility modulating enzymatic activities, signaling pathways, and protein-protein or protein-lipid interactions [82,83]. Since lipid rafts are known entry platforms for many viruses, and sterol/glycosphingolipid depletion drastically impairs infection in other systems [84]. We therefore suggest that sterol oxidation, trafficking, and remodelling contribute to antiviral defense in *O. mediterraneus* by altering plasma membrane organisation and protein-lipid interactions, ultimately influencing resistance or susceptibility.

### Conclusions

This study provides the first comprehensive characterization of two immune phenotypes in *O. mediterraneus*, revealing dynamic lipid remodelling and the potential role of oxidized sterols and lipids in antiviral defense, while reinforcing the central role of the SOC in shared defense mechanisms across *Ostreococcus* species. Our findings reveal intricate metabolic reprogramming in R and S lines, with integrated transcriptomic and metabolomic analyses uncovering congruent patterns in lipid biosynthetic pathways, though most differentially expressed genes were not directly linked to phenotype-specific metabolite biomarkers. Methodological limitations, such as incomplete metabolomic coverage (*e.g.*, glycosylation pathways in detergent-insoluble microdomains), as well as the high variability of the metabolomic profiles, likely contributed to discrepancies between transcriptomic and metabolomic signatures, highlighting the need for broader analytical approaches. Future high-frequency sampling during viral infection, viral lipidomic profiling, and functional analyses—will be essential to fully resolve the temporal and regulatory cascades underlying host-virus interactions in marine phytoplankton.

## Materials and Methods

### Culture conditions

The virus resistant strain (R line) used in this experiment refers to a subculture of *Ostreococcus mediterraneus* RCC2590 (also known as ML line), while the susceptible strain (S line, also known as MA3 and S3), derived from the R strain through single cell dilution method [11,85]. The S strain had been obtained during experimental evolution of the R strain and spontaneously developed susceptibility to Ostreococcus mediterraneus Virus 2 (OmV2). Cultures were grown in a Keller Artificial Seawater medium (K-ASWO) containing 0.42 M NaCl (Sigma S3014), 10 mM KCl (Sigma P9333), 20 mM MgCl_2_ (Sigma M2670), 10 mM CaCl_2_ (Sigma C5080), 24.5 mM MgSO_4_ (Sigma M5921), 2.5 mM NaHCO_3_ (Sigma S6297), complemented with the following Keller stock solutions (NCMA Bigelow) added at 1 mL per liter of medium. The final concentrations in the medium were as follows: 882 µM NaNO_3_; 50.1 µM NH_4_Cl; 10 µM Na_2_ β-glycerophosphate; 10 nM H_2_SeO_3_; 1 mM Tris-HCl (pH 7.2). Trace metals were present at final concentrations of 112 µM Na_2_EDTA, 11.7 µM FeCl_3_, 26 nM Na_2_MoO_4_, 76.5 nM ZnSO_4_, 42 nM CoCl_2_, 910 nM MnCl_2_, 39.2 nM CuSO_4_. Vitamins were supplied at final concentrations of 0.369 nM cyanocobalamin, 2.05 nM biotin and 296 nM thiamine [86–89]. *O. mediterraneus* strains were cultured in T75 cell culture flasks with a ventilated cap (Sarstedt, Germany) containing 200 ml of K-ASWO medium. For routine culture maintenance, 200 µl of culture was transferred into 200 ml of fresh K-ASWO culture medium every two to three weeks. Cultures were maintained at 20°C under a 12:12 h light/dark cycle (100 µE m^-2^ s^-1^) and manually agitated once per day.

To monitor growth kinetics, each flask was inoculated at an initial cell density of 1.00×10^5^ cells.ml^-1^. Cell density was measured daily by flow cytometry on a subset of five representative flasks for each of the R and S strains. (Table S14, Figure S4).

### Culture axenisation

To minimise bacterial contamination, all cultures were treated with a combination of antibiotics. Specifically, 50 µg/mL ampicillin (A9518, Sigma-Aldrich), 50 µg/mL gentamycin (G1914, Sigma-Aldrich), 20 µg/mL kanamycin (60615, Sigma-Aldrich), and 100 µg/mL neomycin (N6386, Sigma-Aldrich) were added to the K-ASWO medium. Following two rounds of subculturing, the bacterial load was reduced to less than 1%, enabling the subsequent transcriptomic and metabolomic analyses (Table S15).

### Experimental design

To investigate transcriptomic and metabolomic changes between the resistant (R) and susceptible (S) strains, we sampled each strain at four time points over 24 hours (Figure S1). Cells were harvested eight days post inoculation at an initial cell density of 1.00×10^5^ cells.ml^-1^ while in exponential growth (Figure S4) Time points *t_1_* and *t_4_,* as well as *t_2_* and *t_3_,* correspond to 2 hours after and 2 hours before the light and dark phases, respectively. Additionally, *t_2_* and *t_3_* mark the beginning and end of the *Ostreococcus* cell division cycle [90]. Metabolomic analyses were performed only at *t_1_* and *t_3_,* whereas transcriptomic analyses were performed at all four time points. Thus, for each time point, phenotype and analysis type, three independent culture flasks of 200 mL were used as biological replicates, resulting in a total of 36 flasks (Figure S1).

### Flow cytometry

Cells were fixed with glutaraldehyde (0.25% final concentration; G6257, Sigma-Aldrich) and Pluronic F-68 (0.1% final concentration; P-7061, Sigma-Aldrich) for 15 minutes in the dark [91]. Following fixation, cells were stained with SYBR Green I (LON50512, Ozyme) for additional 15 minutes in the dark. Cell enumeration was performed using a Beckman-Coulter Cytoflex flow cytometer (laser excitation wavelength: 488 nm). Picoalgae were identified by chlorophyll autofluorescence (detection filter: > 620 nm), while bacteria were detected based on SYBR Green I fluorescence (detection bandwidth: 525-540 nm, corresponding to the FITC channel) and side scatter. Data analysis was conducted using CytExpert 2.2 software (Beckman-Coulter).

### Transcriptomic sampling, RNA extraction and sequencing

For each of the four time points, three replicates of 100 mL cultures of S and R were collected, and cells were harvested by centrifugation at 8000 *g* for 10 min at 20°C. Each cell pellet was resuspended into 2 mL sterile seawater in a 2 mL Eppendorf tube and recentrifuged for another 4 min (8000 *g*, 20°C). The cell pellet was retained, flash frozen in liquid nitrogen and stored at - 80°C until RNA extraction. Total RNA was extracted using the Direct-zol RNA MiniPrep kit (R2052, Zymo Research) and assessed for quality with a Bioanalyzer 2100 (Agilent). Selection for polyadenylated RNA, library preparation and sequencing were performed four times per sample at the Bio-environnement platform (Perpignan University, France). Total RNA samples were concentrated to 1000 ng and treated with NEB Next Ultra Directional RNA kit (New England BioLabs), then each library was amplified through 10 PCR cycles. All 96 libraries (2 strains x 4 timepoints x 3 biological replicates x 4 technical replicates) were sequenced on a NextSeq550 system (Illumina) by multiplexing all samples on a single HighOutput flow cell lane, which generating paired end reads of 101 bp in length. RNA sequence reads were checked for quality using FastQC v0.12.1 (https://www.bioinformatics.babraham.ac.uk/projects/fastqc/).

### Differential gene transcription analysis

Transcriptome read pairs (fragments) were first trimmed to remove sequence adaptors using TrimGalore v0.6.10 (https://www.bioinformatics.babraham.ac.uk/projects/trim_galore/). Index and read alignment onto the annotated *O. mediterraneus* reference genome sequence, including the nuclear, the mitochondrial and the chloroplastic genomes, as well as the OmV2 genome, was performed using STAR v2.7.10b [92]. The raw counts of fragments aligning to each gene were determined using RSEM v1.3.3 [93]. Scripts are available in supplementary material. The raw read counts were first explored by PCA, using gene expression data after applying a multi-step normalisation procedure designed to minimise technical biases and highlight biologically meaningful variation. Only genes with at least 10 mapped fragments in at least one sample were retained. First, raw read counts were normalised by gene length to account for transcript size. Second, counts were scaled by the total number of reads per library to correct for differences in sequencing depth across samples. Third, expression values for each gene were normalised by their total expression across all samples to emphasise relative temporal dynamics rather than absolute abundance. The resulting expression matrix was then centred and scaled (z-score normalisation) prior to PCA. PCA was performed in the R v4.2.1 statistical environment, with transcriptomes (n = 24) as individuals and normalised gene expression values as variables. Genes contributing most strongly to the fourth principal component (PC4), which separates resistant and susceptible phenotypes, were identified and compared to the list of genes involved in the biosynthetic pathways leading to biomarker metabolites.

Differential host gene expression analyses were performed on raw count table (Table S16) using the R package DESeq2 v1.26 [94]. Gene transcription was compared between resistant and susceptible strains at each time point. Genes with a *p*_adj_ < 0.01 were considered significantly differentially transcribed between strains. Furthermore, only genes with a Log_2_ Fold Change (LFC) greater than 2 or lower than -2, corresponding to a minimum fourfold increase or decrease in expression levels, were retained. Genes were considered transcribed if at least 10 reads were mapped to the gene.

### Functional analysis of differentially expressed genes

Functional enrichment analysis was carried out using the Pico-Plaza platform [95](https://bioinformatics.psb.ugent.be/plaza/versions/plaza_pico_03/) to identify overrepresented functional categories among the differentially expressed genes (DEGs). Analyses were performed separately for Gene Ontology (GO) terms, covering biological process, cellular component, and molecular function, and for InterPro domains. For both types of annotations, only protein-coding genes were considered, and enrichment statistics were computed exclusively on genes with at least one GO annotation. A Bonferroni correction was applied to adjust for multiple testing, to retain GO terms or InterPro domains with a *p*_value_ < 0.05. For GO analyses, enrichment settings included GO data from all available sources and no filtering based on evidence codes. The background model used for all enrichment tests corresponded to the annotated gene set of *Ostreococcus mediterraneus*. To account for temporal variation in gene expression, functional enrichment analyses were conducted independently for each time point. Additionally, DEGs on the SOC chromosome, suspected to play a key role in immunity and possibly harbour biomarkers for susceptibility or resistance, were analysed separately from those on other chromosomes. Particular attention was focused on DEGs that were differentially expressed between phenotypes at all time points, as these likely represent stable molecular signatures of resistance or susceptibility. For genes lacking GO or InterPro annotation, either located on the SOC or systematically differentially expressed across all four time points on standard chromosomes, protein domain predictions were examined to infer potential functions based on conserved architectures.

### Orthology analyses between SOC genes of *O. tauri* and *O. mediterraneus*

To assess the conservation of Small Outlier Chromosome (SOC) genes between *Ostreococcus tauri* and *Ostreococcus mediterraneus*, we performed a reciprocal best hit (RBH) analysis. The analysis was based on the complete set of annotated protein-coding genes from both species, retrieved from the ORCAE database (https://bioinformatics.psb.ugent.be/orcae/). Prior to alignment, low-complexity regions were masked using the -no_dust option in BLAST to avoid spurious matches. No e-value threshold was applied, to maximise sensitivity for detecting highly diverged orthologs. Only genes that were reciprocal best hits between the two species were considered orthologous.

### Metabolite extraction and UHPLC-HRMS analyses

At time point *t_1_* and *t_3_*, R and S cultures were harvested by filtration of 100 ml of culture through a Whatman GF/F filter (Z242519, Sigma-Aldrich) under reduced pressure (600 mbar). The filters were subsequently placed into disposable glass culture tubes, ground with 7 ml of ethyl acetate (16371, Sigma-Aldrich), and the solvent was immediately evaporated. The resulting dry extracts were stored at -80°C prior to analysis.

The extracts were analysed as described by Marcellin-Gros et al. (2020). Briefly, an Ultimate 3000 UHPLC Dionex system coupled to an Orbitrap MS/MS FT Q-Exactive Focus Thermo Scientific mass spectrometer was used. Samples were solubilised in MeOH (1 mg. mL^-1^) and 1 µL was injected onto a Phenomenex Luna Omega Polar C18 column (150 × 2.1 mm, 1.6 µm, 100 Å), conditioned at 42°C. The mobile phase consisted of water (solvent A) and acetonitrile (solvent B, 012041, Biosolve), both modified with 0.1% formic acid. The gradient was as follows: 50% B from 3 min before injection to 1 min after, a linear increase of B to 85% from 1 to 3 min, 85% B for 2 min, 89% B from 5.1 to 7 min, 93% B from 7.1 to 10 min, 97% B from 10.1 to 13 min, and 100% B from 13.1 to 18 min. The flow rate was set to 0.5 mL.min-1, with analysis performed 1 min after injection. Mass spectrometry was conducted in positive electrospray ionisation mode, with a mass range of 133.4-2000 Da and centroid mode for mass spectra. FullMS was performed with a resolution of 70,000, AGC target set to 3.0×10^6^ for a chromatogram peak width (FWHM) of 6 s. MS2 had a resolution of 17,500, AGC target of 1.0×10^5^, isolation windows of 0.4 Da, and stepped normalised collision energy of 15/30/45, with dynamic exclusion for 10 s. Lock mass calibration was performed on the Cu(CH_3_CN)^2+^ ion at m/z 144.9821 Da.

### LC-MS data pre-processing

Total ion chromatograms (TIC) were processed using the lipidomic workflow of MS-Dial v4.16 software [96]. A Quality Control mix (QC), composed of the twelve picoalgal extracts, was analysed together with the picoalgal extracts and K-ASWO medium, which served as a blank to remove non-picoalgal compounds. The MS-Dial workflow performs retention time correction, detection of unknown compounds, and grouping across samples, fills gaps when features are absent, removes chemical background using blank samples, and predicts the elemental composition of compounds. The retention time window was set to 2-17 min, with a maximum time shift for compound alignment of 0.1 min. The maximum mass tolerance for compound grouping and elemental composition calculation was 3 ppm, and the minimum peak intensity was set to 1.0×10^6^. This workflow generated an observation/variable matrix, which was used for subsequent statistical analysis.

### Identification of R and S associated metabolites

To identify metabolites associated with either the susceptible or the resistant phenotype, comparative analysis was performed using the S/R ratio. Student’s *t*-tests were applied on feature relative abundances under the null hypothesis of equal ion abundance between strains. Metabolites with *p*_value_ below 0.05 were considered significantly differentially abundant, and an additional fold change threshold of 2 (equivalent to 1 in log_2_ scale) was applied to retain only the most pronounced differences. Additionally, all ions identified as significant were manually inspected to eliminate duplicates corresponding to the same metabolite and to remove false positives. Metabolite identification was carried out by comparing MS^2^ fragmentation spectrum with Sirius 5.6.2 software [26] and raw molecular formulas, calculated from high-resolution mass spectrometry, against databases such as the Dictionary of Marine Natural Products (https://dnp.chemnetbase.com) and SciFinder (American Chemical Society) to retrieve candidate natural compounds.

### Identification of biosynthesis and catabolic pathways of metabolites biomarkers

The metabolic pathways investigated here were reconstructed by integrating multiple complementary sources of information. Genes were initially identified in *Ostreococcus tauri* using the KEGG database (https://www.genome.jp/kegg/), and their orthologues or homologous in *O. mediterraneus* were subsequently retrieved through BLAST searches in Pico-Plaza. Gene and protein domain annotations were further refined using ORCAE, while characterised orthologues in *A. thaliana* were retrieved from TAIR (https://www.arabidopsis.org/). In addition, relevant literature and domain-based searches (InterPro and Pfam) were used to complement the annotation process, ensuring a comprehensive identification of the gene repertoire underlying these pathways.

## Acknowledgments

We would like to thank the Genotoul Bioinformatics platform Toulouse Occitanie (Bioinfo Genotoul, doi: 10.15454/1.5572369328961167E12) for providing computing and storage resources. We are grateful to the Platform BioPIC and Bio2mar platforms for the access to the cytometry and UHPLC-HRMS equipment, respectively, which are supported by EMBRC-France and AO-EMBRC, whose French state funds are managed by the ANR within the Investments of the Future program under reference ANR-10-INBS-0 and ANR-21-ESRE-0038. We thank all past and present GENOPHY group members for both stimulation discussions and technical support.

## Data availability

The reference genome is available on Genbank under accession number GCA_012295225.1. The annotations have been downloaded from ORCAE (https://bioinformatics.psb.ugent.be/gdb/O.mediterraneus/). The 24 transcriptomes generated for this study have been submitted to ENA under Project number #.

The metabolomics data has been submitted to #.

## List of Supplemental Figures

**Figure S1.**
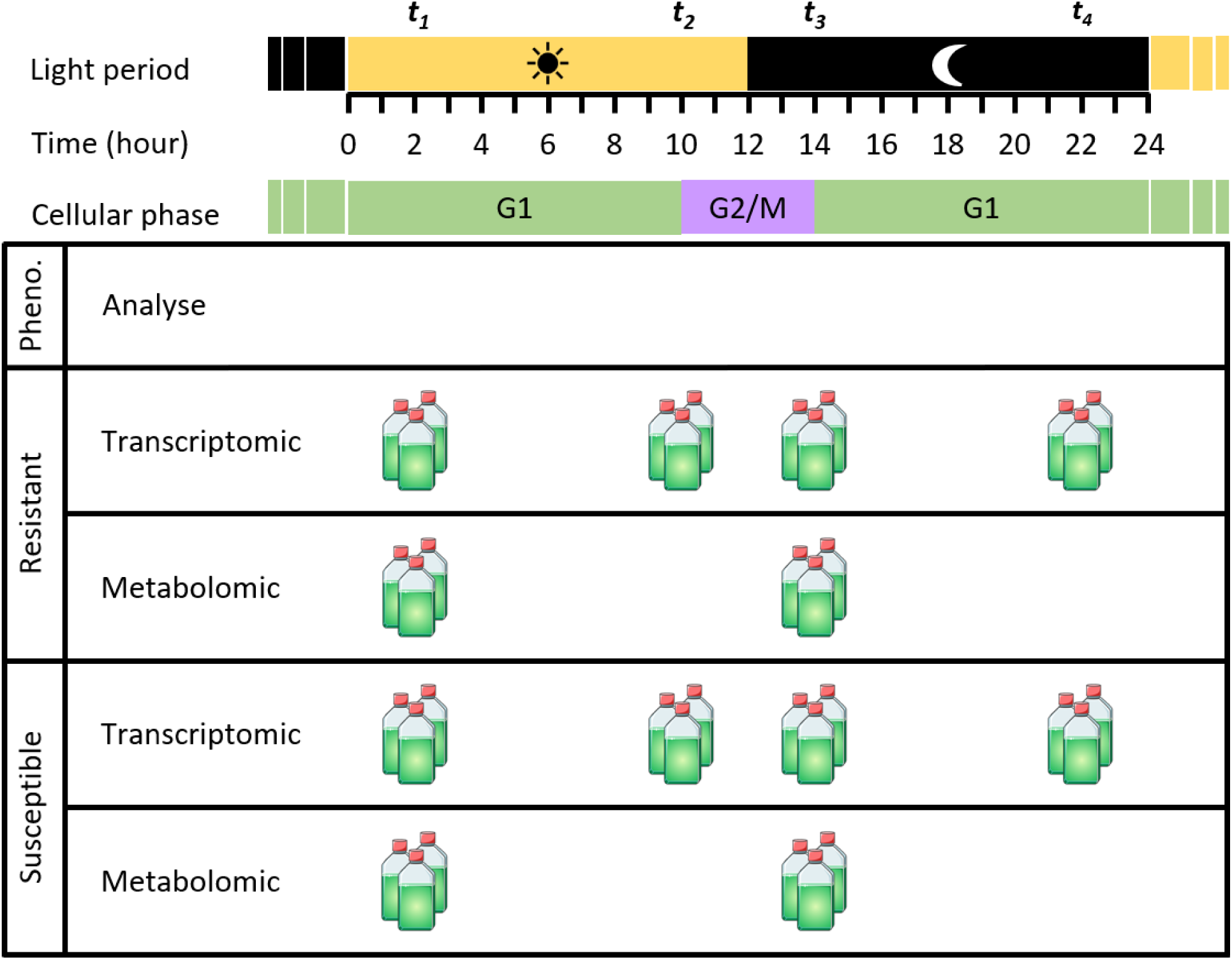
Experimental design for the transcriptomic and metabolomic analysis of *Ostreococcus mediterraneus* at *t_1_*, *t_2_*, *t_3_*, and *t_4_*. G1: cell growth; G2/M: preparation for division/mitosis.

**Figure S2.**
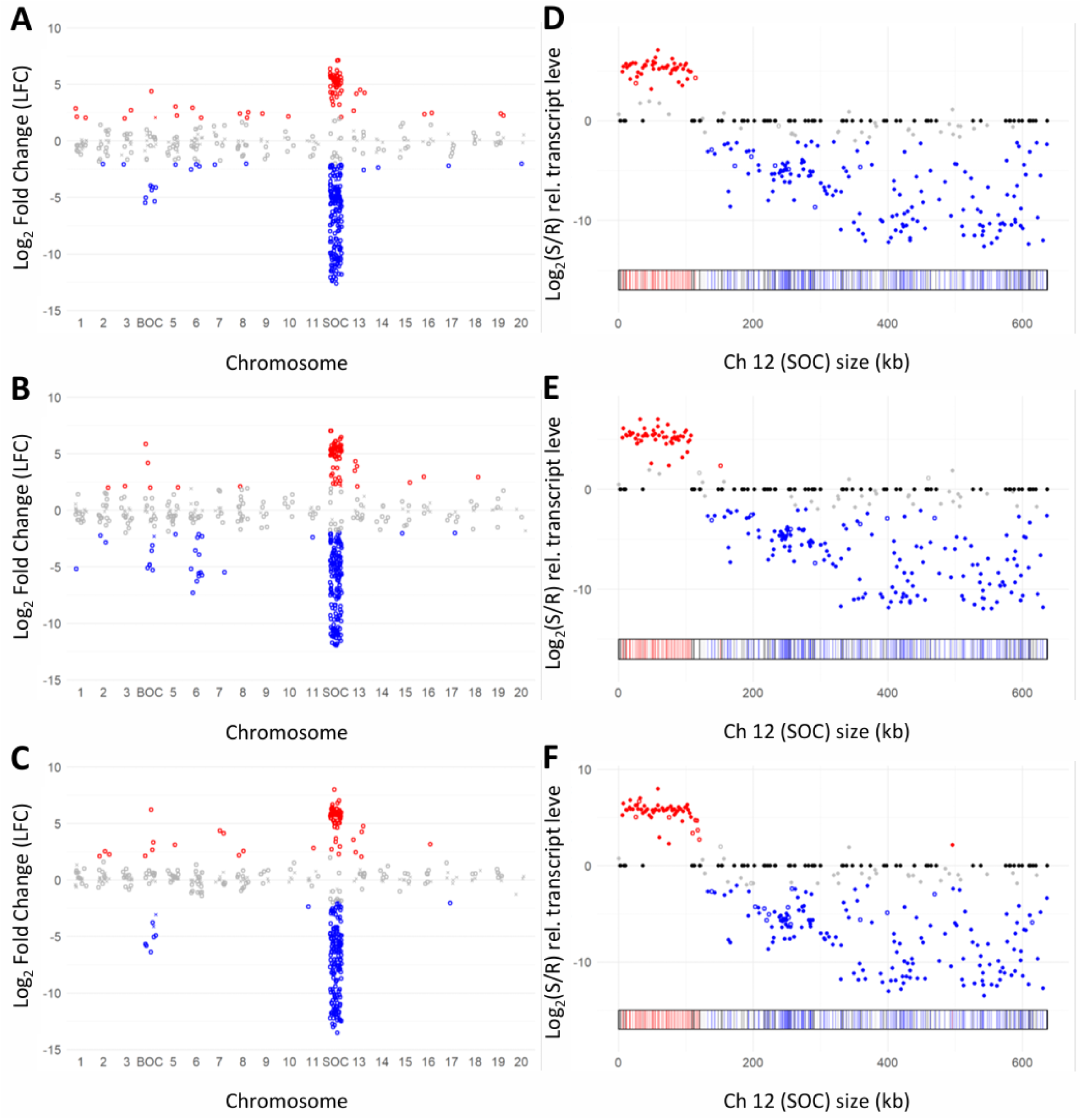
Differential gene expression across and within chromosomes. **Left part:** Distribution of log_2_ fold change (LFC) values across chromosomes for 477 differentially expressed genes (DEGs) at **A**) *t_2_*, **B**) *t_3_* and **C**) *t_4_*. Genes overexpressed (|LFC| ≥ 2) in the S line are shown in red and in the R line in blue, while values below this threshold are grey. Significant values are shown as open circles and non-significant ones as crosses. **Right part:** Bipartite distribution of gene expression on the SOC at **D**) *t_2_*, **E**) *t_3_* and **F**) *t_4_*. Genes overexpressed in the S line are shown in red, those overexpressed in the R line in blue, not differentially expressed genes in grey, and non-expressed genes in black. Filled dots represent On/Off genes, while empty dots indicate genes that are not On/Off.

**Figure S3.**
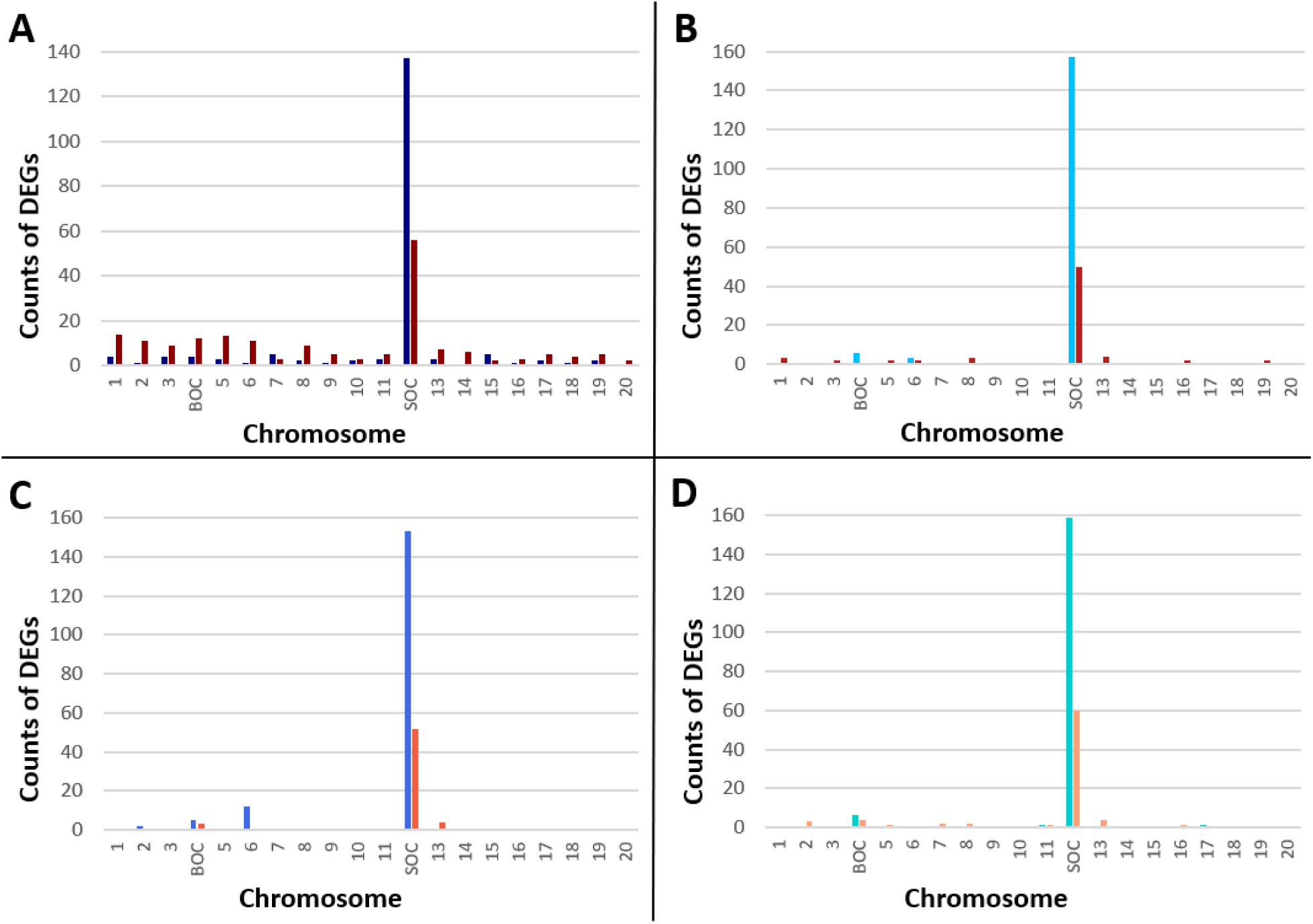
Histograms showing the number of overexpressed genes per chromosome in each line across time points: **A**) *t_1_*, **B**) *t_2_*, **C**) *t_3_*, and **D**) *t_4_*.

**Figure S4.**
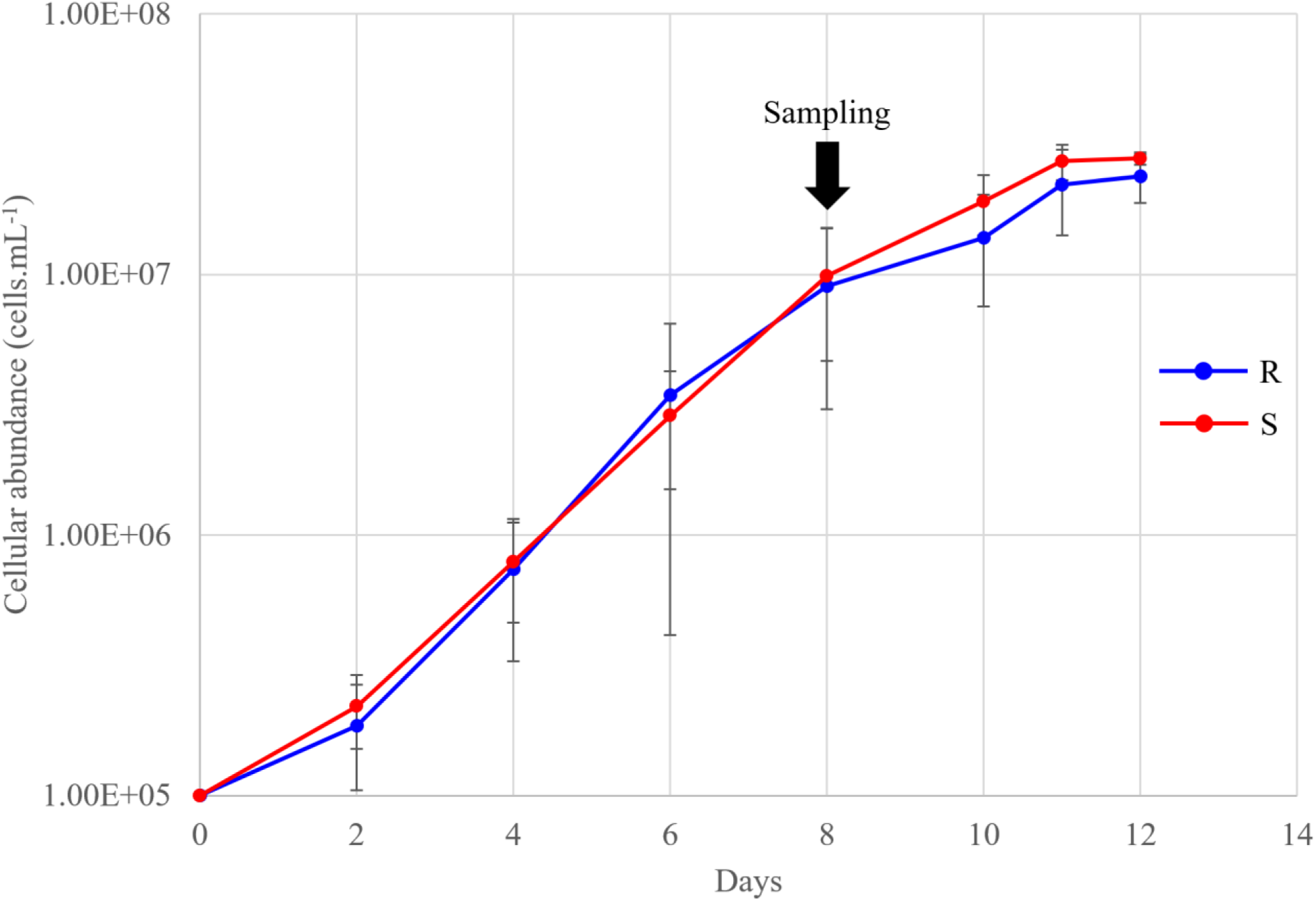
Growth kinetics of the R (blue) and S (red) strain of *Ostreococcus mediterraneus* (12/12 light/dark cycle, 20°C)

## References

1. Mojica KDA, Brussaard CPD. Marine Viruses and Their Role in Marine Ecosystems and Carbon Cycling. Ann Rev Mar Sci. 2026;18: 351–380. doi:10.1146/annurev-marine-040324-020244

2. Courties C, Vaquer A, Troussellier M, Lautier J, Chrétiennot-Dinet MJ, Neveux J, et al. Smallest eukaryotic organism. Nature. 1994;370: 255–255. doi:10.1038/370255a0

3. Derelle E, Ferraz C, Rombauts S, Rouzé P, Worden AZ, Robbens S, et al. Genome analysis of the smallest free-living eukaryote *Ostreococcus tauri* unveils many unique features. Proceedings of the National Academy of Sciences. 2006;103: 11647–11652. doi:10.1073/pnas.0604795103

4. Jancek S, Gourbiere S, Moreau H, Piganeau G. Clues about the Genetic Basis of Adaptation Emerge from Comparing the Proteomes of Two Ostreococcus Ecotypes (Chlorophyta, Prasinophyceae). Mol Biol Evol. 2008;25: 2293–2300. doi:10.1093/molbev/msn168

5. Tragin M, Vaulot D. Novel diversity within marine Mamiellophyceae (Chlorophyta) unveiled by metabarcoding. Sci Rep. 2019;9: 5190. doi:10.1038/s41598-019-41680-6

6. Schulz F, Roux S, Paez-Espino D, Jungbluth S, Walsh DA, Denef VJ, et al. Giant virus diversity and host interactions through global metagenomics. Nature. 2020;578: 432–436. doi:10.1038/s41586-020-1957-x

7. Clerissi C, Desdevises Y, Grimsley N. Prasinoviruses of the Marine Green Alga Ostreococcus tauri Are Mainly Species Specific. J Virol. 2012;86: 4611–4619. doi:10.1128/JVI.07221-11

8. Yung CCM, Rey Redondo E, Sanchez F, Yau S, Piganeau G. Diversity and Evolution of Mamiellophyceae: Early-Diverging Phytoplanktonic Green Algae Containing Many Cosmopolitan Species. J Mar Sci Eng. 2022;10: 240. doi:10.3390/jmse10020240

9. Baudoux A-C., Lebredonchel H, Dehmer H, Latimier M, Edern R, Rigaut-Jalabert F, et al. Interplay between the genetic clades of *Micromonas* and their viruses in the Western English Channel. Environ Microbiol Rep. 2015;7: 765–773. doi:10.1111/1758-2229.12309

10. Wilhelm SW, Suttle CA. Viruses and Nutrient Cycles in the Sea. Bioscience. 1999;49: 781–788. doi:10.2307/1313569

11. Yau S, Krasovec M, Benites LF, Rombauts S, Groussin M, Vancaester E, et al. Virus-host coexistence in phytoplankton through the genomic lens. Sci Adv. 2020;6. doi:10.1126/sciadv.aay2587

12. Subirana L, Péquin B, Michely S, Escande M-L, Meilland J, Derelle E, et al. Morphology, Genome Plasticity, and Phylogeny in the Genus Ostreococcus Reveal a Cryptic Species, O. mediterraneus sp. nov. (Mamiellales, Mamiellophyceae). Protist. 2013;164: 643–659. doi:10.1016/j.protis.2013.06.002

13. Blanc-Mathieu R, Krasovec M, Hebrard M, Yau S, Desgranges E, Martin J, et al. Population genomics of picophytoplankton unveils novel chromosome hypervariability. Sci Adv. 2017;3. doi:10.1126/sciadv.1700239

14. Yau S, Hemon C, Derelle E, Moreau H, Piganeau G, Grimsley N. A Viral Immunity Chromosome in the Marine Picoeukaryote, Ostreococcus tauri. PLoS Pathog. 2016;12: e1005965. doi:10.1371/journal.ppat.1005965

15. Rosenwasser S, Mausz MA, Schatz D, Sheyn U, Malitsky S, Aharoni A, et al. Rewiring Host Lipid Metabolism by Large Viruses Determines the Fate of *Emiliania huxleyi*, a Bloom-Forming Alga in the Ocean. Plant Cell. 2014;26: 2689–2707. doi:10.1105/tpc.114.125641

16. Vardi A, Van Mooy BAS, Fredricks HF, Popendorf KJ, Ossolinski JE, Haramaty L, et al. Viral Glycosphingolipids Induce Lytic Infection and Cell Death in Marine Phytoplankton. Science (1979). 2009;326: 861–865. doi:10.1126/science.1177322

17. Schleyer G, Shahaf N, Ziv C, Dong Y, Meoded RA, Helfrich EJN, et al. In plaque-mass spectrometry imaging of a bloom-forming alga during viral infection reveals a metabolic shift towards odd-chain fatty acid lipids. Nat Microbiol. 2019;4: 527–538. doi:10.1038/s41564-018-0336-y

18. Ziv C, Malitsky S, Othman A, Ben-Dor S, Wei Y, Zheng S, et al. Viral serine palmitoyltransferase induces metabolic switch in sphingolipid biosynthesis and is required for infection of a marine alga. Proceedings of the National Academy of Sciences. 2016;113. doi:10.1073/pnas.1523168113

19. Zhang E, Gao J, Wei Z, Zeng J, Li J, Li G, et al. MicroRNA-mediated regulation of lipid metabolism in virus-infected *Emiliania huxleyi*. ISME J. 2022;16: 2457–2466. doi:10.1038/s41396-022-01291-y

20. Schleyer G, Kuhlisch C, Ziv C, Ben-Dor S, Malitsky S, Schatz D, et al. Lipid biomarkers for algal resistance to viral infection in the ocean. Proceedings of the National Academy of Sciences. 2023;120. doi:10.1073/pnas.2217121120

21. Keeling PJ, Burki F, Wilcox HM, Allam B, Allen EE, Amaral-Zettler LA, et al. The Marine Microbial Eukaryote Transcriptome Sequencing Project (MMETSP): Illuminating the Functional Diversity of Eukaryotic Life in the Oceans through Transcriptome Sequencing. PLoS Biol. 2014;12: e1001889. doi:10.1371/journal.pbio.1001889

22. Romero-Losada AB, Arvanitidou C, García-Gómez ME, Morales-Pineda M, Castro-Pérez MJ, Chew YP, et al. Multiomics integration unveils photoperiodic plasticity in the molecular rhythms of marine phytoplankton. Plant Cell. 2025;37. doi:10.1093/plcell/koaf033

23. Monnier A, Liverani S, Bouvet R, Jesson B, Smith JQ, Mosser J, et al. Orchestrated transcription of biological processes in the marine picoeukaryote Ostreococcus exposed to light/dark cycles. BMC Genomics. 2010;11: 192. doi:10.1186/1471-2164-11-192

24. Pietrangelo A, Ridgway ND. Bridging the molecular and biological functions of the oxysterol-binding protein family. Cellular and Molecular Life Sciences. 2018;75: 3079–3098. doi:10.1007/s00018-018-2795-y

25. Marcellin-Gros R, Piganeau G, Stien D. Metabolomic Insights into Marine Phytoplankton Diversity. Mar Drugs. 2020;18: 78. doi:10.3390/md18020078

26. Dührkop K, Fleischauer M, Ludwig M, Aksenov AA, Melnik A V., Meusel M, et al. SIRIUS 4: a rapid tool for turning tandem mass spectra into metabolite structure information. Nat Methods. 2019;16: 299–302. doi:10.1038/s41592-019-0344-8

27. Guella G, Frassanito R, Mancini I. A new solution for an old problem: the regiochemical distribution of the acyl chains in galactolipids can be established by electrospray ionization tandem mass spectrometry. Rapid Communications in Mass Spectrometry. 2003;17: 1982–1994. doi:10.1002/rcm.1142

28. Mimouni V, Couzinet-Mossion A, Ulmann L, Wielgosz-Collin G. Lipids From Microalgae. Microalgae in Health and Disease Prevention. Elsevier; 2018. pp. 109–131. doi:10.1016/B978-0-12-811405-6.00005-0

29. Rivera SM, Christou P, Canela-Garayoa R. Identification of carotenoids using mass spectrometry. Mass Spectrom Rev. 2014;33: 353–372. doi:10.1002/mas.21390

30. Domergue F, Abbadi A, Zähringer U, Moreau H, Heinz E. *In vivo* characterization of the first acyl-CoA Δ6-desaturase from a member of the plant kingdom, the microalga *Ostreococcus tauri*. Biochemical Journal. 2005;389: 483–490. doi:10.1042/BJ20050111

31. Ahmann K, Heilmann M, Feussner I. Identification of a Δ4-desaturase from the microalga *Ostreococcus lucimarinus*. European Journal of Lipid Science and Technology. 2011;113: 832–840. doi:10.1002/ejlt.201100069

32. Tavares S, Grotkjær T, Obsen T, Haslam RP, Napier JA, Gunnarsson N. Metabolic Engineering of *Saccharomyces* cerevisiae for Production of Eicosapentaenoic Acid, Using a Novel Δ5-Desaturase from Paramecium tetraurelia. Appl Environ Microbiol. 2011;77: 1854–1861. doi:10.1128/AEM.01935-10

33. Degraeve-Guilbault C, Gomez RE, Lemoigne C, Pankansem N, Morin S, Tuphile K, et al. Plastidic Δ6 Fatty-Acid Desaturases with Distinctive Substrate Specificity Regulate the Pool of C18-PUFAs in the Ancestral Picoalga *Ostreococcus tauri*. Plant Physiol. 2020;184: 82–96. doi:10.1104/pp.20.00281

34. Estévez JM, Cantero A, Reindl A, Reichler S, León P. 1-Deoxy-d-xylulose-5-phosphate Synthase, a Limiting Enzyme for Plastidic Isoprenoid Biosynthesis in Plants. Journal of Biological Chemistry. 2001;276: 22901–22909. doi:10.1074/jbc.M100854200

35. Marshall B, Amritkar K, Wolfe M, Kaçar B, Landick R. Evolutionary flexibility and rigidity in the bacterial methylerythritol phosphate (MEP) pathway. Front Microbiol. 2023;14. doi:10.3389/fmicb.2023.1286626

36. Worden AZ, Lee J-H, Mock T, Rouzé P, Simmons MP, Aerts AL, et al. Green Evolution and Dynamic Adaptations Revealed by Genomes of the Marine Picoeukaryotes *Micromonas*. Science (1979). 2009;324: 268–272. doi:10.1126/science.1167222

37. Moreau H, Verhelst B, Couloux A, Derelle E, Rombauts S, Grimsley N, et al. Gene functionalities and genome structure in Bathycoccus prasinos reflect cellular specializations at the base of the green lineage. Genome Biol. 2012;13: R74. doi:10.1186/gb-2012-13-8-r74

38. Bugnot C, Alioto T, Rodriguez FC, Garrido JG, Gut M, Yau S, et al. Chromoanagenesis is a driver of structural variation in the smallest photosynthetic eukaryote. 2025. doi:10.1101/2025.09.17.676765

39. Bojarová P, Williams SJ. Sulfotransferases, sulfatases and formylglycine-generating enzymes: a sulfation fascination. Curr Opin Chem Biol. 2008;12: 573–581. doi:10.1016/j.cbpa.2008.06.018

40. Henquet M, Lehle L, Schreuder M, Rouwendal G, Molthoff J, Helsper J, et al. Identification of the Gene Encoding the α1,3-Mannosyltransferase (ALG3) in *Arabidopsis* and Characterization of Downstream *N*-Glycan Processing. Plant Cell. 2008;20: 1652–1664. doi:10.1105/tpc.108.060731

41. Schoberer J, Strasser R. Sub-Compartmental Organization of Golgi-Resident N-Glycan Processing Enzymes in Plants. Mol Plant. 2011;4: 220–228. doi:10.1093/mp/ssq082

42. Kang BS, Baek JH, Macoy DM, Chakraborty R, Cha J-Y, Hwang D-J, et al. N-Glycosylation process in both ER and Golgi plays pivotal role in plant immunity. Journal of Plant Biology. 2015;58: 374–382. doi:10.1007/s12374-015-0197-3

43. Li J, Zhao-Hui C, Batoux M, Nekrasov V, Roux M, Chinchilla D, et al. Specific ER quality control components required for biogenesis of the plant innate immune receptor EFR. Proceedings of the National Academy of Sciences. 2009;106: 15973–15978. doi:10.1073/pnas.0905532106

44. Chen X-L, Shi T, Yang J, Shi W, Gao X, Chen D, et al. *N*-Glycosylation of Effector Proteins by an α-1,3-Mannosyltransferase Is Required for the Rice Blast Fungus to Evade Host Innate Immunity. Plant Cell. 2014;26: 1360–1376. doi:10.1105/tpc.114.123588

45. Pontejo SM, Murphy PM. Mouse Cytomegalovirus Differentially Exploits Cell Surface Glycosaminoglycans in a Cell Type-Dependent and MCK-2-Independent Manner. Viruses. 2019;12: 31. doi:10.3390/v12010031

46. Salvador B, Sexton NR, Carrion R, Nunneley J, Patterson JL, Steffen I, et al. Filoviruses Utilize Glycosaminoglycans for Their Attachment to Target Cells. J Virol. 2013;87: 3295–3304. doi:10.1128/JVI.01621-12

47. McAllister N, Liu Y, Silva LM, Lentscher AJ, Chai W, Wu N, et al. Chikungunya Virus Strains from Each Genetic Clade Bind Sulfated Glycosaminoglycans as Attachment Factors. J Virol. 2020;94. doi:10.1128/JVI.01500-20

48. Cohen M. Notable Aspects of Glycan-Protein Interactions. Biomolecules. 2015;5: 2056–2072. doi:10.3390/biom5032056

49. Piacente F, Gaglianone M, Laugieri M, Tonetti M. The Autonomous Glycosylation of Large DNA Viruses. Int J Mol Sci. 2015;16: 29315–29328. doi:10.3390/ijms161226169

50. Tanaka R, Oster U, Kruse E, Rüdiger W, Grimm B. Reduced Activity of Geranylgeranyl Reductase Leads to Loss of Chlorophyll and Tocopherol and to Partially Geranylgeranylated Chlorophyll in Transgenic Tobacco Plants Expressing Antisense RNA for Geranylgeranyl Reductase1. Plant Physiol. 1999;120: 695–704. doi:10.1104/pp.120.3.695

51. Li G, Woroch AD, Donaher NA, Cockshutt AM, Campbell DA. A Hard Day’s Night: Diatoms Continue Recycling Photosystem II in the Dark. Front Mar Sci. 2016;3. doi:10.3389/fmars.2016.00218

52. Derelle E, Yau S, Moreau H, Grimsley NH. Prasinovirus Attack of Ostreococcus Is Furtive by Day but Savage by Night. J Virol. 2018;92. doi:10.1128/JVI.01703-17

53. Jian W, Zhang D, Zhu F, Wang S, Zhu T, Pu X, et al. Nitrate reductase-dependent nitric oxide production is required for regulation alternative oxidase pathway involved in the resistance to Cucumber mosaic virus infection in Arabidopsis. Plant Growth Regul. 2015;77: 99–107. doi:10.1007/s10725-015-0040-3

54. Sheyn U, Rosenwasser S, Ben-Dor S, Porat Z, Vardi A. Modulation of host ROS metabolism is essential for viral infection of a bloom-forming coccolithophore in the ocean. ISME J. 2016;10: 1742–1754. doi:10.1038/ismej.2015.228

55. Britto DT, Kronzucker HJ. Cellular mechanisms of potassium transport in plants. Physiol Plant. 2008;133: 637–650. doi:10.1111/j.1399-3054.2008.01067.x

56. García-Martí M, Piñero MC, García-Sanchez F, Mestre TC, López-Delacalle M, Martínez V, et al. Amelioration of the Oxidative Stress Generated by Simple or Combined Abiotic Stress through the K+ and Ca2+ Supplementation in Tomato Plants. Antioxidants. 2019;8: 81. doi:10.3390/antiox8040081

57. Hirth M, Liverani S, Mahlow S, Bouget F-Y, Pohnert G, Sasso S. Metabolic profiling identifies trehalose as an abundant and diurnally fluctuating metabolite in the microalga Ostreococcus tauri. Metabolomics. 2017;13: 68. doi:10.1007/s11306-017-1203-1

58. Monnier A, Liverani S, Bouvet R, Jesson B, Smith JQ, Mosser J, et al. Orchestrated transcription of biological processes in the marine picoeukaryote Ostreococcus exposed to light/dark cycles. BMC Genomics. 2010;11: 192. doi:10.1186/1471-2164-11-192

59. Degraeve-Guilbault C, Bréhélin C, Haslam R, Sayanova O, Marie-Luce G, Jouhet J, et al. Glycerolipid Characterization and Nutrient Deprivation-Associated Changes in the Green Picoalga *Ostreococcus tauri*. Plant Physiol. 2017;173: 2060–2080. doi:10.1104/pp.16.01467

60. Buseman CM, Tamura P, Sparks AA, Baughman EJ, Maatta S, Zhao J, et al. Wounding Stimulates the Accumulation of Glycerolipids Containing Oxophytodienoic Acid and Dinor-Oxophytodienoic Acid in Arabidopsis Leaves. Plant Physiol. 2006;142: 28–39. doi:10.1104/pp.106.082115

61. Andersson MX, Hamberg M, Kourtchenko O, Brunnström Å, McPhail KL, Gerwick WH, et al. Oxylipin Profiling of the Hypersensitive Response in Arabidopsis thaliana. Journal of Biological Chemistry. 2006;281: 31528–31537. doi:10.1016/S0021-9258(19)84066-8

62. Vu HS, Tamura P, Galeva NA, Chaturvedi R, Roth MR, Williams TD, et al. Direct Infusion Mass Spectrometry of Oxylipin-Containing Arabidopsis Membrane Lipids Reveals Varied Patterns in Different Stress Responses. Plant Physiol. 2012;158: 324–339. doi:10.1104/pp.111.190280

63. Kourtchenko O, Andersson MX, Hamberg M, Brunnström A, Göbel C, McPhail KL, et al. Oxo-Phytodienoic Acid-Containing Galactolipids in Arabidopsis: Jasmonate Signaling Dependence. Plant Physiol. 2007;145: 1658–1669. doi:10.1104/pp.107.104752

64. Sattler SE, Mène-Saffrané L, Farmer EE, Krischke M, Mueller MJ, DellaPenna D. Nonenzymatic Lipid Peroxidation Reprograms Gene Expression and Activates Defense Markers in *Arabidopsis* Tocopherol-Deficient Mutants. Plant Cell. 2006;18: 3706–3720. doi:10.1105/tpc.106.044065

65. Zoeller M, Stingl N, Krischke M, Fekete A, Waller F, Berger S, et al. Lipid Profiling of the Arabidopsis Hypersensitive Response Reveals Specific Lipid Peroxidation and Fragmentation Processes: Biogenesis of Pimelic and Azelaic Acid. Plant Physiol. 2012;160: 365–378. doi:10.1104/pp.112.202846

66. Zhang P, Jackson E, Li X, Zhang Y. Salicylic acid and jasmonic acid in plant immunity. Hortic Res. 2025;12. doi:10.1093/hr/uhaf082

67. Wasternack C, Strnad M. Jasmonate signaling in plant stress responses and development – active and inactive compounds. N Biotechnol. 2016;33: 604–613. doi:10.1016/j.nbt.2015.11.001

68. Stintzi A, Weber H, Reymond P, Browse J, Farmer EE. Plant defense in the absence of jasmonic acid: The role of cyclopentenones. Proceedings of the National Academy of Sciences. 2001;98: 12837–12842. doi:10.1073/pnas.211311098

69. Schäfer M, Fischer C, Meldau S, Seebald E, Oelmüller R, Baldwin IT. Lipase Activity in Insect Oral Secretions Mediates Defense Responses in Arabidopsis. Plant Physiol. 2011;156: 1520–1534. doi:10.1104/pp.111.173567

70. Arnold MD, Gruber C, Floková K, Miersch O, Strnad M, Novák O, et al. The Recently Identified Isoleucine Conjugate of cis-12-Oxo-Phytodienoic Acid Is Partially Active in cis-12-Oxo-Phytodienoic Acid-Specific Gene Expression of Arabidopsis thaliana. PLoS One. 2016;11: e0162829. doi:10.1371/journal.pone.0162829

71. Figueiredo A, Martins J, Sebastiana M, Guerreiro A, Silva A, Matos AR, et al. Specific adjustments in grapevine leaf proteome discriminating resistant and susceptible grapevine genotypes to Plasmopara viticola. J Proteomics. 2017;152: 48–57. doi:10.1016/j.jprot.2016.10.012

72. Laureano G, Figueiredo J, Cavaco AR, Duarte B, Caçador I, Malhó R, et al. The interplay between membrane lipids and phospholipase A family members in grapevine resistance against Plasmopara viticola. Sci Rep. 2018;8: 14538. doi:10.1038/s41598-018-32559-z

73. Hartmann M. Plant sterols and the membrane environment. Trends Plant Sci. 1998;3: 170–175. doi:10.1016/S1360-1385(98)01233-3

74. Clouse SD. Arabidopsis Mutants Reveal Multiple Roles for Sterols in Plant Development. Plant Cell. 2002;14: 1995–2000. doi:10.1105/tpc.140930

75. Ketter E, Randall G. Virus Impact on Lipids and Membranes. Annu Rev Virol. 2019;6: 319–340. doi:10.1146/annurev-virology-092818-015748

76. Mackenzie JM, Khromykh AA, Parton RG. Cholesterol Manipulation by West Nile Virus Perturbs the Cellular Immune Response. Cell Host Microbe. 2007;2: 229–239. doi:10.1016/j.chom.2007.09.003

77. Blanc M, Hsieh WY, Robertson KA, Watterson S, Shui G, Lacaze P, et al. Host Defense against Viral Infection Involves Interferon Mediated Down-Regulation of Sterol Biosynthesis. PLoS Biol. 2011;9: e1000598. doi:10.1371/journal.pbio.1000598

78. Raychaudhuri S, Im YJ, Hurley JH, Prinz WA. Nonvesicular sterol movement from plasma membrane to ER requires oxysterol-binding protein–related proteins and phosphoinositides. J Cell Biol. 2006;173: 107–119. doi:10.1083/jcb.200510084

79. Umate P. Oxysterol binding proteins (OSBPs) and their encoding genes in Arabidopsis and rice. Steroids. 2011;76: 524–529. doi:10.1016/j.steroids.2011.01.007

80. Laloi M, Perret A-M, Chatre L, Melser S, Cantrel C, Vaultier M-N, et al. Insights into the Role of Specific Lipids in the Formation and Delivery of Lipid Microdomains to the Plasma Membrane of Plant Cells. Plant Physiol. 2007;143: 461–472. doi:10.1104/pp.106.091496

81. Simon-Plas F, Perraki A, Bayer E, Gerbeau-Pissot P, Mongrand S. An update on plant membrane rafts. Curr Opin Plant Biol. 2011;14: 642–649. doi:10.1016/j.pbi.2011.08.003

82. Schaller H. The role of sterols in plant growth and development. Prog Lipid Res. 2003;42: 163–175. doi:10.1016/S0163-7827(02)00047-4

83. Valitova JN, Sulkarnayeva AG, Minibayeva F V. Plant sterols: Diversity, biosynthesis, and physiological functions. Biochemistry (Moscow). 2016;81: 819–834. doi:10.1134/S0006297916080046

84. Phalen T, Kielian M. Cholesterol is required for infection by Semliki Forest virus. J Cell Biol. 1991;112: 615–623. doi:10.1083/jcb.112.4.615

85. Krasovec M, Eyre-Walker A, Sanchez-Ferandin S, Piganeau G. Spontaneous Mutation Rate in the Smallest Photosynthetic Eukaryotes. Mol Biol Evol. 2017;34: 1770–1779. doi:10.1093/molbev/msx119

86. Sambrook Joseph, Russell DW. Molecular cloning: a laboratory manual. Cold Spring Harbor Laboratory Press; 2001.

87. Keller MD, Selvin RC, Claus W, Guillard RRL. Media for the culture of oceanic ultraphytoplankton. J Phycol. 1987;23: 633–638. doi:10.1111/j.1529-8817.1987.tb04217.x

88. Guillard RRL. Culture of Phytoplankton for Feeding Marine Invertebrates. Culture of Marine Invertebrate Animals. Boston, MA: Springer US; 1975. pp. 29–60. doi:10.1007/978-1-4615-8714-9_3

89. Guillard RRL, Ryther JH. Studies of marine planktonic diatoms. I. Cyclotella nana Hustedt, and Detonula confervacea (cleve) Gran. Can J Microbiol. 1962;8: 229–239. doi:10.1139/m62-029

90. Farinas B, Mary C, de O Manes C-L, Bhaud Y, Peaucellier G, Moreau H. Natural Synchronisation for the Study of Cell Division in the Green Unicellular Alga Ostreococcus tauri. Plant Mol Biol. 2006;60: 277–292. doi:10.1007/s11103-005-4066-1

91. Marie D, Rigaut-Jalabert F, Vaulot D. An improved protocol for flow cytometry analysis of phytoplankton cultures and natural samples. Cytometry Part A. 2014;85: 962–968. doi:10.1002/cyto.a.22517

92. Dobin A, Davis CA, Schlesinger F, Drenkow J, Zaleski C, Jha S, et al. STAR: ultrafast universal RNA-seq aligner. Bioinformatics. 2013;29: 15–21. doi:10.1093/bioinformatics/bts635

93. Li B, Dewey CN. RSEM: accurate transcript quantification from RNA-Seq data with or without a reference genome. BMC Bioinformatics. 2011;12: 323. doi:10.1186/1471-2105-12-323

94. Love MI, Huber W, Anders S. Moderated estimation of fold change and dispersion for RNA-seq data with DESeq2. Genome Biol. 2014;15: 550. doi:10.1186/s13059-014-0550-8

95. Vandepoele K, Van Bel M, Richard G, Van Landeghem S, Verhelst B, Moreau H, et al. pico-PLAZA, a genome database of microbial photosynthetic eukaryotes. Environ Microbiol. 2013;15: 2147–2153. doi:10.1111/1462-2920.12174

96. Tsugawa H, Ikeda K, Takahashi M, Satoh A, Mori Y, Uchino H, et al. A lipidome atlas in MS-DIAL 4. Nat Biotechnol. 2020;38: 1159–1163. doi:10.1038/s41587-020-0531-2

